# Pyrethroid exposure alters *Anopheles albimanus* microbiota and resistant mosquitoes harbor more insecticide-metabolizing bacteria

**DOI:** 10.1101/537480

**Authors:** Nsa Dada, Juan C. Lol, Ana Cristina Benedict, Francisco López, Mili Sheth, Nicole Dzuris, Norma Padilla, Audrey Lenhart

## Abstract

A deeper understanding of the mechanisms underlying insecticide resistance is needed to mitigate its threat to malaria vector control. Building upon our earlier identified associations between mosquito microbiota and insecticide resistance, we demonstrate for the first time, type-specific effects of pyrethroid exposure on internal and cuticle surface bacteria in adult progeny of field-collected *Anopheles albimanus*. In contrast, larval cuticle surface—but not internal—bacteria were affected by pyrethroid exposure. Being over five-folds more abundant in pyrethroid resistant adults, as compared to susceptible or non-insecticide-exposed mosquitoes, *Klebsiella* (alphacypermethrin), *Pantoea* and *Asaia* (permethrin) were identified as potential markers of pyrethroid resistance in *An. albimanus*. We also show for the first time that *An. albimanus* larvae and adult cuticles harbor more diverse bacterial communities than their internal microbial niches. Our findings indicate insecticide selection pressures on mosquito microbiota, and support the hypothesis of an undescribed microbe-mediated mechanism of insecticide metabolism in mosquitoes.

**Competing interest statement:** This work was supported by the US Centers for Disease Control and Prevention (CDC) through the American Society for Microbiology’s (ASM) Infectious Disease and Public Health Microbiology Postdoctoral Fellowship program, and the CDC’s Advanced Molecular Detection (AMD) program. The findings and conclusions in this paper are those of the authors and do not necessarily represent the official position of the CDC or ASM.

## Introduction

Recent evidence suggests that progress in global malaria control has stalled, with an estimated 2 million more malaria cases in 2017 than in 2016, and an increase in malaria incidence in the region of the Americas (*1*). This stall in progress overlaps with increasing reports of insecticide resistance (*1, 2*), which poses a growing challenge to malaria vector control programs (*3*). The mechanisms underlying insecticide resistance in malaria vectors in the Americas remain poorly characterized, despite intensified vector control efforts stemming from regional malaria elimination programs (*4*).

Four main mechanisms of insecticide resistance have been described in mosquitoes (*5*): cuticular modifications, insecticide target site insensitivity, heightened insecticide detoxification and behavioral avoidance of insecticides. While these aforementioned mechanisms have been described in malaria vectors across multiple geographical regions (*5*), there remain significant gaps in our understanding of the mechanisms of insecticide resistance and resistance intensity in mosquito populations. However, increasing access to advanced genomic tools has made it easier to investigate other aspects of mosquito biology, such as their microbiota, that may be associated with insecticide resistance.

Mosquitoes, like other living organisms, are hosts to a variety of microbes that are principally acquired from their breeding habitats during immature development, and from adult food sources (*6*). In addition to habitat and/or food source-acquired microbes, transstadial bacterial transmission from adult females to their eggs (*7*), across immature stages (*8*) and unto the adult stage (*7*) have also been demonstrated in mosquitoes. These microbes, some of which are known to metabolize insecticides (*9-11*), actively shape their host physiology (*6, 12*). We postulated that the mosquito microbiota may thus contribute to insecticide detoxification, consequently augmenting resistance in the host; a phenomenon that has previously been demonstrated in agricultural pests (*13*), but only now being investigated in disease vectors (*10, 14*). We previously studied the microbiota of field-caught Peruvian *Anopheles albimanus* (*10*) and detected significant differences in the composition and putative functions of bacteria between fenitrothion-resistant and -susceptible mosquitoes. These results provided a comprehensive baseline regarding the bacterial composition of *An. albimanus* in relation to insecticide resistance, and suggested that the mosquito microbiota could be impacted by insecticides, and/or contributing to insecticide resistance.

Malaria vector control programs in the Americas rely heavily on pyrethroid insecticides, used mainly on bednets or for indoor residual spraying (IRS) (*15*), thus increasing pyrethroid selection pressure on malaria vector populations. However, in this region, information on the underlying mechanisms of pyrethroid resistance in malaria vectors is scarce. To elucidate these mechanisms of pyrethroid resistance, especially from the perspective of the microbiota, the present study focuses on elucidating the effects of pyrethroid exposure on the microbiota of *An. albimanus* from Guatemala. Our objectives were to characterize and compare the composition of internal and cuticle surface bacteria between adult and larval F1 progeny of field-collected mosquitoes that were either exposed (further classified as pyrethroid-resistant or –susceptible), or not exposed to pyrethroid insecticides. In order to check for spatial consistency, we tested F1 progeny originating from multiple locations. We discuss our findings in light of the hypothesis that mosquito microbiota is impacted by insecticides, and contribute to the detoxification of insecticides within the host. We also report the first comprehensive characterization of *An. albimanus* larval microbiota, as well as the mosquito cuticle surface microbiota.

## Materials and Methods

### Mosquito collections and mass rearing

Gravid adult female *An. albimanus* were collected in and around four cattle corrals sampled in two villages: El Terrero and Las Cruces, in La Gomera, Escuintla, Guatemala (Fig. 1). Mosquito samples, separated by collection site, were held in paper cups with access to cotton pads soaked in 10% sucrose solution, and transported to the insectary at Universidad del Valle de Guatemala in Guatemala City for species identification and oviposition. Mosquitoes were morphologically sorted using identification keys (*16*). Approximately 300 gravid females identified as *An. albimanus* based on the presence of a white terminal palpal segment, white third and fourth hind tarsomeres, and basal dark band on fifth hindtarsomere, were used for oviposition. Species identification was confirmed by PCR (as described below) performed on a subsample of the F_1_ progeny. Oviposition was achieved using a previously described method (*17*) with modifications. Briefly, ovipostion containers (ovipots) were created by placing gravid females in 32 oz. paper soup containers (60±10 mosquitoes/container) containing distilled water to a depth of 2-3 cm. Prior to the introduction of mosquitoes, the containers were covered with netting, subsequently topped with cotton pads soaked in 10% sucrose solution, then covered with a thick black plastic bag to trap in moisture and keep the containers dark. The ovipots were kept under standard insectary conditions of 27±2°C, 80±10% relative humidity and a 12 h light-dark cycle for at least 48 hours to allow for ovipostion, after which adults were removed and eggs collected. Under the aforementioned conditions, immature mosquitoes were reared using distilled water, with F_1_ progeny of parents from the same collection site grouped together and reared separately from those of parents from other sites. Eggs from ovipots were washed into 18 x 14 x 3 inch plastic larval trays (150-200 eggs per tray) containing distilled water to a depth of 2-3 cm, with 3-4 drops of 10% yeast solution. Hatched larvae were fed finely ground Koi food (Foster & Smith, Inc. Rhinelander, WI) until pupation. Approximately a quarter of the proportion of resulting third to fourth instar larvae (L3-L4) were collected for insecticide resistance assays, and the remaining larvae were reared until the pupal stage. With the aid of a stereo microscope, male and female pupae were separated into 8 oz. paper cups placed in cardboard cages for adult emergence. Adult virgin females were provided 10% sucrose solution until they were 2-5 days old and were then used for insecticide resistance assays.

**Fig. 1.**
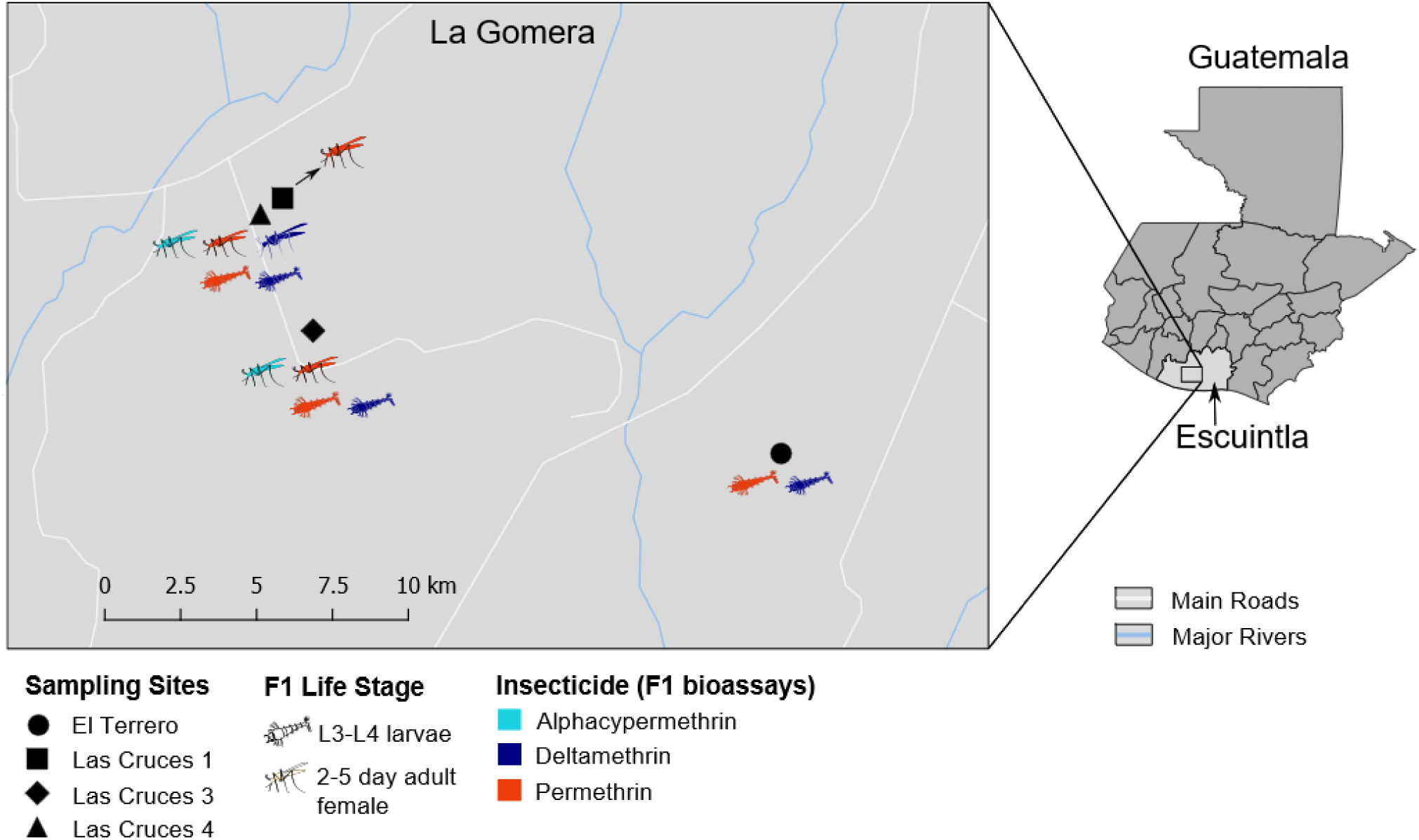
Sampling sites of field-caught *An. albimanus* and details of bioassays on resulting F_1_ progeny. This figure shows the map of Guatemala indicating the Department of Escuintla (right), and an expanded view of La Gomera Municipality indicating the four sites from which field-collected mosquitoes were sampled (left). F_1_ Larvae (n =132) and adult (n =135) progeny originating from these sites were tested for insecticide resistance, and for each collection site, the figure shows the developmental stage of F_1_ progeny tested, color-coded by the type of insecticide they were tested for. See Suppl. 2 for a summary of the number of larvae and adults processed for each insecticide per site.

### Determination of pyrethroid resistance status in larval and adult F_1_ progeny of field-collected *An. albimanus*

Three different pyrethroid insecticides were used for this study: alphacypermethrin, deltamethrin and permethrin. Adult mosquitoes were tested for resistance to all three insecticides, while larvae were tested with only deltamethrin and permethrin (Fig. 1). Prior to determining the pyrethroid resistance status of the F_1_ L3-L4 larvae from field-caught mosquitoes, the diagnostic dose (a dose that kills 100% of susceptible mosquitoes within a given time) and diagnostic time (the expected time to achieve 100% mortality) (*18*) for each insecticide were determined (Suppl. 1A) using L3-L4 larvae from the insecticide-susceptible *An. albimanus* Sanarate strain (*19*). Insecticide resistance assays were performed in the insectary under the aforementioned rearing conditions. For larvae, 100 ml of distilled water was added to 150 ml glass beakers, followed by 1 ml of insecticide stock solution. Absolute ethanol without added insecticide was added to the control beaker. The beakers were left to stand for 10-15 mins to allow the solvent to evaporate. Subsequently, 15-20 L3-L4 larvae were added to each beaker, and mortality scored at the end of the diagnostic time (Suppl. 1A). Larvae that were alive at the end of the assays were classified as resistant, those that were dead or moribund were classified as susceptible, while those that were used in the control beaker (all survived) were classified as non-exposed. For each insecticide tested, three replicates of the assay were run, and resistant, susceptible, and non-exposed samples from all replicates were pooled for further processing.

The adult insecticide resistance assays were performed on 2-5 day old non-blood fed virgin F_1_ females using the CDC bottle bioassay method as previously described (*18*). Each insecticide was tested using the recommended diagnostic dose and time (*18*) for *Anopheles* mosquitoes (Suppl. 1B), and at the end of the assays, mosquitoes were classified as resistant, susceptible or non-exposed as above. For both larvae and adults, resistant and non-exposed samples were euthanized by placing them in liquid nitrogen or at - 80°C. All samples were preserved in RNALater® (Applied Biosystems, Foster City, CA) and held in - 80°C until shipping (on dry ice) to the US Centers for Disease Control and Prevention (CDC) in Atlanta, USA, where the molecular analyses were performed.

### Extraction and purification of genomic DNA from whole larvae and adult mosquitoes and the surface of their cuticles

A total of 132 larvae and 135 adults that had been categorized as resistant, susceptible or non-exposed to each tested insecticide were processed. Prior to genomic DNA extraction, mosquito samples were thawed at 4°C overnight, and the RNALater® solution was subsequently discarded. Samples were then rinsed once with nuclease free water to remove any remaining RNALater® solution and pooled for further processing. Samples were grouped by parental collection site, and for each insecticide tested, three pools of mosquitoes (at both the larval and adult stages), comprising three mosquitoes per pool, were processed in each category unless otherwise stated. To obtain the microbiota from the surface of the cuticle, each pool of larvae or adult mosquitoes was washed in nuclease free water by agitating with a vortex mixer for 15-20 seconds, and the resulting wash solution was transferred to a new tube for DNA extraction. The pools of whole larvae or adults were subsequently surface sterilized as previously described (*10*).

Genomic DNA was extracted from pools of surface sterilized whole larvae and adult *An. albimanus* as well as from the washed cuticle surface of each respective pool, using the DNeasy Blood and Tissue Kit (QIAGEN, Hilden, Germany) with slight modifications. Each pool was homogenized in 180 µL buffer ATL (QIAGEN) using the TissueLyser II (QIAGEN) with two sets of 96-well adapter plates (QIAGEN) containing 2 ml microcentrifuge tubes (QIAGEN) in which samples were placed, each along with 5 mm diameter stainless steel beads (QIAGEN). The TissueLyser II (QIAGEN) was set to 30 hz/sec for a total of 15 mins for the pools of whole mosquito samples, and eight minutes for the washed cuticle samples, with plates rotated every minute until the end of the homogenization process. Homogenized samples were transferred to new tubes for further processing. Twenty micro liters of Proteinase K (QIAGEN) was added to each homogenized sample and incubated at 56°C overnight, following which 200 µL of buffer AL (QIAGEN) was added and incubated for an additional 2 hours at the same temperature. At the end of the incubation period, 200 µL of 100% ethanol was added to the mixture and transferred to spin columns (QIAGEN). The remaining wash steps were performed following QIAGEN’s spin-column protocol for the purification of total DNA from animal tissues, and the purified DNA was eluted in 70 µL buffer AE (QIAGEN). Two negative (sans samples) extraction controls each were processed along with the internal and cuticle surface microbiota samples. All steps (as well as those described below) were performed under sterile conditions. Genomic DNA was stored at −80°C until further processing.

### PCR for mosquito species identification and 16s rRNA gene amplification

Amplification of the second internal transcribed spacer region (ITS2) of the ribosomal DNA was used to confirm the morphological identification of *An. albimanus*. This was performed on DNA from individual mosquito legs that had been dissected prior to washing the cuticle surface, from a subsample of individual adult mosquitoes. The universal ITS2 primers (ITS2 A: TGTGAACTGCAGGACACAT and ITS2 B: TATGCTTAAATTCAGGGGGT) for distinguishing members of cryptic *Anopheles* complexes (*20*) was used in a PCR reaction volume of 25 µL, comprising ≥100 ng/µL DNA template, 15 µM of each respective primer, 12.5 µL of 2x AccuStart II PCR SuperMix (Quanta Bio, Beverly, MA), and PCR grade water to final volume. Reactions were conducted using a T100™ Thermal Cycler (Bio-Rad, USA) with the following conditions: initial denaturation at 94°C for 4 min, then 35 cycles of 94°C for 30 secs, 53°C for 40 secs and 72°C for 30 secs, followed by a final extension for 10 min at 72°C (*21*). Amplified products (~500 bp fragment) resolved by EtBr-stained agarose gel electrophoresis confirmed all samples as *An. albimanus*.

The v3-v4 region of the 16s rRNA gene was amplified from both internal and cuticle surface microbiota samples, along with the negative extraction controls, using the bacterial and archaeal universal primers 341F (**TCGTCGGCAGCGTCAGATGTGTATAAGAGACAG**CCTACGGGNGGCWGCAG) and 805R (**GTCTCGTGGGCTCGGAGATGTGTATAAGAGACAG**GACTACHVGGGTATCTAATCC) with Illumina® adapters (in bold). The PCR was performed using a 25µL reaction volume comprising ≥20 ng/µL DNA template, 5 µM of each primer, 10 µL of 2x KAPA HiFi HotStart PCR mix (Roche, Switzerland), and PCR grade water to final volume. Three negative PCR controls (PCR grade water substituted for DNA template) were processed along with the microbiota samples. Using the aforementioned thermal cycler, the following PCR program was used; initial denaturation at 95°C for 3 min, then 25 cycles of 95°C, 55°C and 72°C for 30 secs each, followed by a final extension for 5 min at 72°C. Resulting amplification products (~460 bp) were resolved as described above and quantified using a NanoDrop™ spectrophotometer (Thermo Fisher Scientific, Waltham, MA). These were subsequently purified using Agencourt AMPure XP beads (Beckman Coulter Inc., Indianapolis, IN, USA) at 0.7x (internal) or 0.875x (cuticle surface) sample volume and eluted in 40 µL 10mM Tris (pH 8.5). Purified PCR products, along with those from all negative controls—which yielded no bands on agarose gel— were submitted to the CDC Biotechnology Core Facility for library preparation and sequencing.

### Library preparation and 16s rRNA sequencing

Index PCR was performed with 25µL NEBNext Hig-Fidelity 2X PCR master mix (New England Biolabs Inc., Ipswich, MA), containing 5µL of each index primer (Nextera XT Index kit v2 set A, B and D; Illumina, San Diego, CA), 10ul of purified PCR products (0-20ng/µL) as template, and PCR grade water to final volume of 50µL. PCR thermal cycler conditions used were: 98°C for 30 sec, followed by 8 cycles of 98°C for 10 sec, 55°C and 65°C for 30 secs each, followed by a final extension at 65°C for 5 min. The resulting products were cleaned using Agencourt AMPure XP beads (Beckman Coulter Inc., Indianapolis, IN, USA) at 1.2x sample volume and eluted in 25 µL 10mM Tris (pH 8.5). Libraries were analyzed for size and concentration, normalized to 2nM and pooled. For cluster generation, the final 2nM pool was denatured following Illumina guidelines for loading onto flowcells. Sequencing was performed on an Illumina Hiseq machine, using Hiseq 2×250 cycle paired-end sequencing kits. Resulting sequence reads were filtered for read quality, basecalled and demultiplexed using bcl2fastq (v2.19).

### Sequencing reads quality control and filtering

The resulting demultiplexed paired-end sequencing reads (115,250,077 in total, with a maximum length of 250 bp) were imported into the ‘quantitative insights into microbial ecology’ pipeline, Qiime2 v.2017.7.0 (*22*). Further read processing and analysis utilizing Qiime2 were performed using v.2018.2.0 of the pipeline. The ‘divisive amplicon denoising algorithm’ `DADA2` (*23*) plugin in Qiime2 was used to ‘denoise’ sequencing reads. This step filters out noise and corrects errors in marginal sequences, removes chimeric sequences and singletons, merges paired-end reads and finally dereplicates the resulting sequences, resulting in high resolution amplicon sequence variants (ASVs) for downstream analysis. Using the `denoise-paired` command, the `DADA2` options passed were `trunc_len_f: 244, trunc_len_r: 244, and n_reads_learn: 500000`, with all other options left as default. This resulted in 30,956,883 ASVs that were further filtered to remove ASVs associated with the extraction and PCR controls (total 4737). Additionally, ASVs with a minimum frequency of 100 were removed, resulting in 17,225,776 ASVs ranging from 3,277 to 223,222 per sample (Suppl. 2).

### Identification of variables significantly impacting the mosquito microbiota

The variables of interest for this study are presented in Suppl. 3. To identify which covariates impacted the microbiota, a multivariate response linear regression model was fit using the `q2-gneiss` (*24*) plugin as described in the Qiime2 v.2018.2.0 manual (https://docs.qiime2.org/2018.2/tutorials/gneiss). This utilizes a ‘leave-one-variable-out’ approach, where one variable at a time is dropped from the model and the change in R2—a measure of the proportion of variance in a dependent variable that is predicted by an independent variable—is calculated to evaluate the impact of a single covariate on the model. To avoid overfitting the model, `q2-gneiss` performs a 10-fold cross validation and reports a mean square error (MSE) for the model and a prediction accuracy MSE at each iteration of the cross validation process—a lower prediction accuracy MSE indicates the absence of model overfitting. All the variables outlined in Suppl. 3 were included in the model, and any covariate whose exclusion from the model resulted in an R2 change >0 was scored as impacting the microbiota—a threshold of R2 change ≥0.01 (i.e. explaining 10% of the variation in the microbiota) was set for significant impact.

### Bacterial diversity

To assess and compare bacterial diversity between samples, the diversity of ASVs within (alpha diversity) and between (beta diversity) samples were computed and compared. The Shannon alpha diversity index (the number of distinct ASVs along with the similarity of their frequencies within a sample)—was computed in QIIME2 using the `q2-diversity` plugin. As alpha diversity metrics are sensitive to uneven sampling depths, multiple rarefactions were performed prior to computing alpha diversity indices; whereby the number of ASVs per sample were randomly selected (without replacement) at an even depth (Suppl. 4). Alpha diversity matrices were compared between samples using the pair-wise Kruskal-Wallis tests with Benjamini-Hochberg false discovery rate (FDR) corrections for multiple comparisons. The Bray-Curtis dissimilarity beta diversity index—the compositional dissimilarity in ASV counts between samples—was computed in QIIME2 and visualized by Principal Co-ordinates Analysis (PCoA) plots in R using the phyloseq R package (*25*). Also using the `q2-diversity` plugin in QIIME2, the Bray-Curtis dissimilarity matrices were computed using both even sampling depth (as normalized above), as well as unnormalized ASV counts. For both methods, pair-wise comparisons of the resulting matrices between samples were computed using permutational (999 permutations) multivariate analysis of variance (PERMANOVA) with FDR corrections. There were negligible differences in outputs between the methods, thus visualization in R was conducted using Bray-Curtis dissimilarity matrices on unnormalized ASV counts. Significance for the pair-wise comparisons was set to *q*-value (i.e. FDR adjusted *p*-value) <0.05.

### Taxonomic annotation and relative abundance of identified bacterial taxa

Taxonomic annotation of ASVs was performed in QIIME 2 using a pre-trained Naïve Bayes classifier (*26*) and the `q2-feature-classifier` plugin (*27*). Prior to annotation, the classifier was trained on the QIIME-compatible 16S SILVA reference (99% identity) database v.128 (*28*). The reference sequences were trimmed to span the v3-v4 region (425 bp) of the 16S rRNA gene using the `extract-reads` command of the `q2-feature-classifier` plugin. The resulting relative abundance tables of annotated ASVs were exported into R and ggplot2 v.2.2.1 (*29*) was used to generate stacked barplots to visualize the relative abundance of bacterial taxa across samples.

### Identification of possible bacterial markers of pyrethroid resistance

The linear discriminant analysis (LDA) effect size method (LEfSe) v.1.7 (*30*) was used to identify possible bacterial markers of pyrethroid resistance. This method identifies genomic features (ASVs in this case) that characterize the differences between two or more biological conditions—it allows for the identification of differentially abundant features that are consistent with biologically meaningful categories (*30*). Mosquito larvae and adults differ in bacterial composition (*31*), and based on consistent outputs from our linear regression analyses and pair-wise beta diversity comparisons, the data from adult and larvae samples were analyzed separately. Microbiota was categorized either as cuticle surface or internal, within which they were grouped by type of insecticide. The relative abundance tables of annotated ASVs, comprising data by type of insecticide tested, were imported into LEfSe—three tables each for the internal and adult cuticle surface microbiota, and two each for the internal and larval cuticle surface microbiota. The data were imported into LEfSe with insecticide resistance phenotype (Res, Sus, Unexp) entered as the `class` vector (the main condition under investigation), and location as `subclass` (as biologically meaningful groupings within each class), and scaled using LEfSe’s 1,10^6^ data transformation. To identify biomarkers, LEfSe first uses pairwise non-parametric factorial Kruskal-Wallis sum-rank test to detect ASVs with significant differential abundance between classes. Then for biological consistency, it performs pairwise unpaired Wilcoxon rank-sum tests to compare differentially abundant ASVs from the previous step between subclasses (location). The default *p*-value <0.05 was used for both tests, and ASVs with consistent significant differential abundance between the three classes and across all locations were identified as possible biomarkers for pyrethroid resistance. Lastly, the biomarkers were used to build an LDA model from which the relative difference among classes was ranked to obtain the effect size. The LDA model was built using default parameters—at least 30 cycles of bootstrapping, each sampling two-thirds of the data with replacement—and the resulting logarithmic score of the analyses are presented. An LDA score of ≥2.0 was set as the threshold to retain a biomarker. All data analysis outputs were edited using Inkscape (*32*). The raw sequencing reads generated from this project, including those from negative controls, have been deposited in the National Center for Biotechnology Information (NCBI), Sequence Read Archive under the BioProject PRJNA512122.

## Results

### Sample characteristics, 16S rRNA sequencing reads and quality control statistics

Suppl. 2 summarizes the number of samples processed for each insecticide per location, the corresponding number of generated sequencing reads, and the number of reads (represented by the percentage of total sequencing reads generated) used for downstream analysis following quality control and dereplication.

### Variations in bacterial composition associated with microbial niche and type of pyrethroid exposure

To determine which variables were associated with bacterial composition in *An. albimanus*, we fit a multivariate response linear regression model using the entire dataset post quality control and dereplication. The model comprising all variables (Suppl. 3) predicted 51% of the variation in bacterial composition (R2=0.51), with insecticide type (R2=0.01), microbial niche (R2=0.04) and developmental stage (R2=0.24) having the highest association (*p*<0.05). Based on this outcome, new models were fit using adult and larval data separately. The impact of insecticide type (R2=0.06) and microbial niche (R2=0.12) remained in adults (*p*<0.05), with the model explaining 26% of the variation in adult bacterial composition (R2=0.26). However, only the microbial niche (R2=0.11) was significantly associated with larval bacterial composition (*p*<0.05), with 39% of the variation being explained by the model (R2=0.39). Based on these outcomes, the adult and larval data were individually categorized by microbial niche and used separately for downstream analysis.

### Pyrethroid insecticide exposure and type impacted bacterial abundance in *An.albimanus*

Comparison of Shannon diversity indices showed a significant impact of both insecticide type and overall exposure on the abundance and evenness of larval cuticle surface bacteria but not internal bacteria, with non-exposed larvae containing the most abundant and even internal and cuticle surface bacterial communities (Suppl. 5). While insecticide exposure significantly impacted bacterial abundance on larval cuticle surfaces, there was no significant difference in bacterial abundance and evenness between pyrethroid-resistant and –susceptible larvae. Conversely, the abundance and evenness of adult internal bacteria, but not those on the cuticle surface, were significantly impacted by type of insecticide (Suppl. 5), with the internal microbiota of deltamethrin-exposed adults representing the most abundant and even bacterial community. No significant impact of overall insecticide exposure was observed on the abundance and evenness of adult cuticle surface bacteria, and insecticide exposure did not significantly impact the internal bacterial abundance between resistant and susceptible adult mosquitoes (Suppl. 5).

### Insecticide-specific impact of pyrethroid exposure on *An. albimanus* microbiota

Irrespective of the type of pyrethroid exposure, comparisons of Bray-Curtis dissimilarity indices showed significant differences in internal, but not cuticle surface, microbiota between pyrethroid-exposed and non-exposed adults (Suppl. 6). In contrast, there was a significant difference in cuticle surface microbiota between pyrethroid-exposed and non-exposed larvae, which was not evident in the internal microbiota (Suppl. 6). In both adults and larvae, the internal and cuticle surface bacteria did not differ between pyrethroid-resistant and susceptible mosquitoes (Suppl. 6). These results were consistent when each type of insecticide tested was considered, except in the case of deltamethrin, where no difference in bacterial composition was detected between exposed and non-exposed mosquitoes (Suppl. 7). Also, there were significant differences (*q*<0.02) in cuticle surface microbiota between alphacypermtherin-exposed and non-exposed adults (Suppl. 7). Bray-Curtis dissimilarity index comparisons further showed significant differences in bacterial composition between adults and larvae (F=117.4, p=0.001), between internal and cuticle surface microbiota in both larvae (F=39.9, p=0.001) and adult mosquitoes (F=38.7, p=0.001), and between samples exposed to different pyrethroids (Suppl. 8), thus corroborating the results of the linear regression analysis.

For each insecticide tested, and across all sampling locations, visualizations of Bray-Curtis diversity distance matrices showed the microbiota of non-exposed mosquitoes clustering distinctly away from those of pyrethroid-exposed mosquitoes, with a gradient separation between resistant and susceptible mosquitoes. This clustering pattern was more evident for the internal microbiota of both adults (Fig. 2) and larvae (Fig. 3) than for the cuticle surface microbiota.

**Fig. 2.**
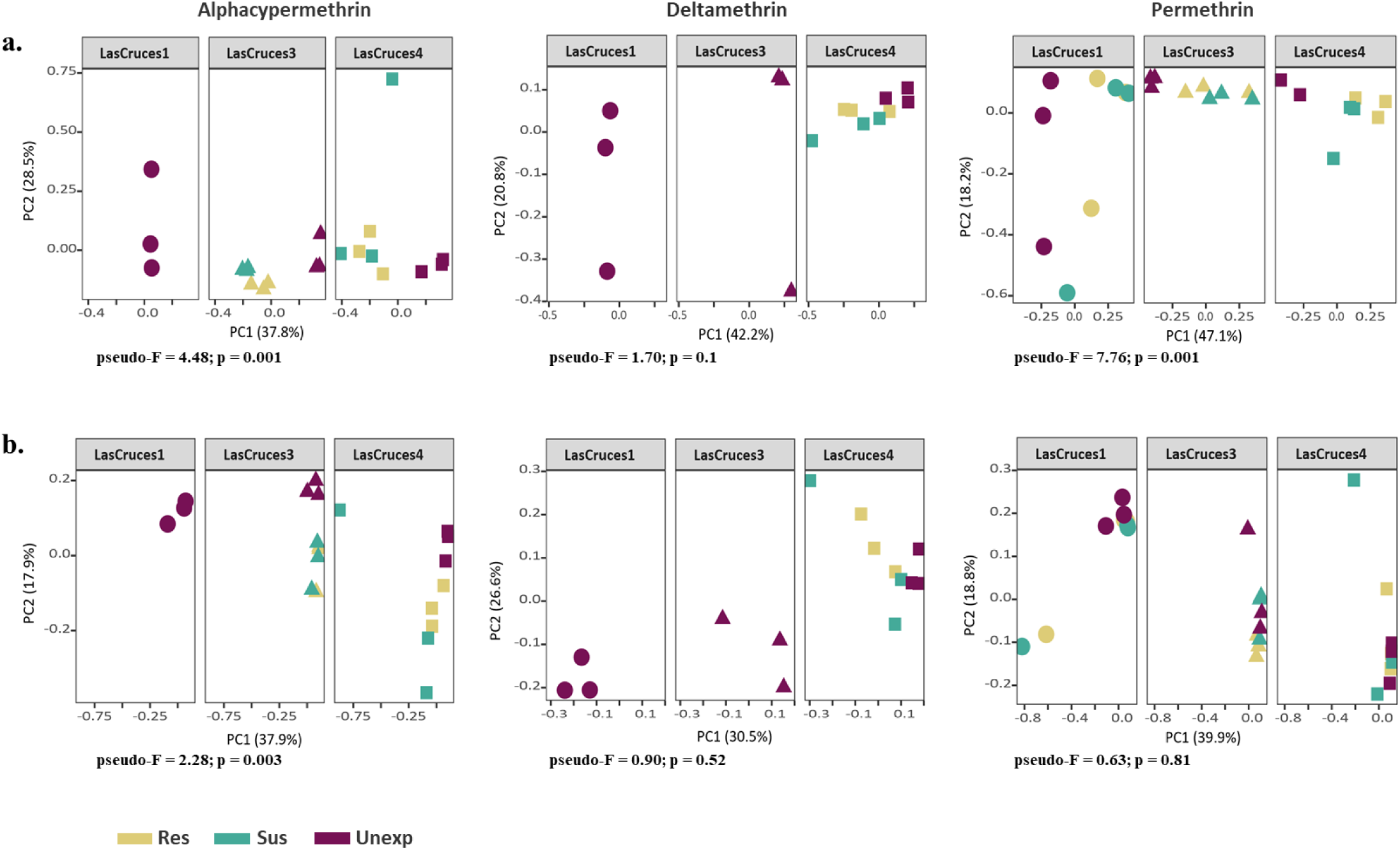
Principal coordinate analysis plots of Bray-Curtis distances between pyrethroid-exposed and non-exposed F_1_ adult *An. albimanus*. The principal coordinate analysis (PCoA) plots, based on Bray-Curtis dissimilarity distances, show clustering patterns of the internal **(A)** and cuticle surface **(B)** microbiota in adult F1 mosquitoes with respect to pyrethroid exposure. Separate plots, further sub-categorized by sample origin (depicted by unique shapes), are presented for each insecticide tested. Each point on the plots represents the bacterial composition of a pool of three mosquitoes, with the axes representing the first two dimensions of the PCoA, along with the proportion (%) of variation in bacterial composition captured. The plots show distinct separation of pyrethroid-exposed (resistant and susceptible) mosquitoes from non-exposed mosquitoes. Within the exposed group, a gradient separation is seen between resistant and susceptible mosquitoes. These clustering patterns are consistent for all three pyrethroid insecticides tested and across all locations for the internal microbiota **(A)**. The cuticle surface microbiota **(B)** however, showed varying clustering patterns with only patterns of alphacypermethrin-tested mosquitoes being consistent with those of the internal microbiota—no distinct clustering patterns were evident in deltamethrin-or permethrin-tested samples. Results of overall non pairwise beta diversity (Bray Curtis) comparison using PERMANOVA (999 permutations) tests are presented. The test statistic value (pseudo-F) for each comparison is presented, with significance set to p-value <0.05. For the internal microbiota **(A)**, the difference in bacterial composition between resistant, susceptible and unexposed mosquitoes was statistically significant for alphacypermethrin and permethrin, whereas in the cuticle surface microbiota **(B)**, this difference was only statistically significant in alphacypermethrin.

**Fig. 3.**
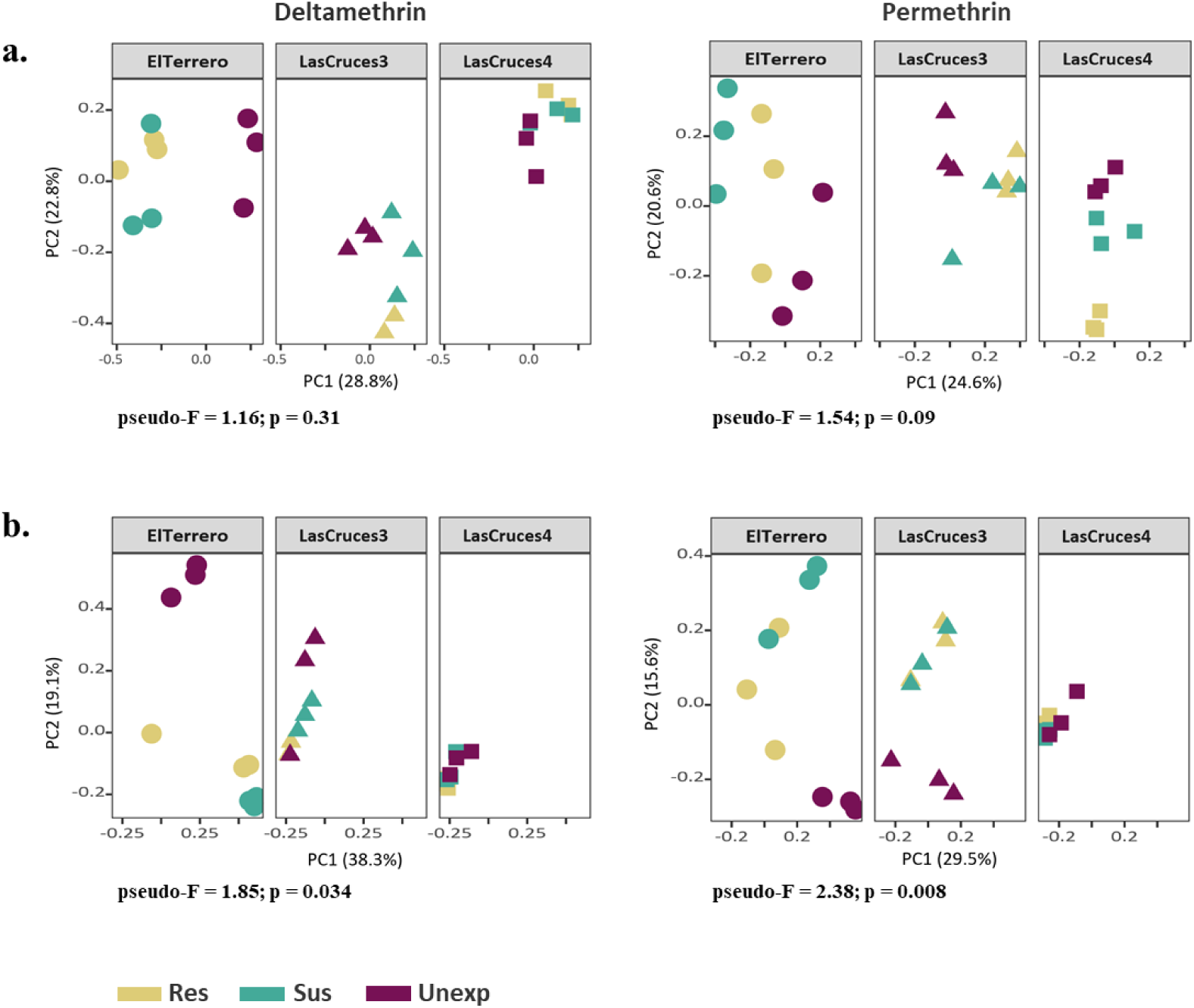
Principal coordinate analysis plots of Bray-Curtis distances between pyrethroid-exposed and non-exposed *An. albimanus* F_1_ larvae. The principal coordinate analysis (PCoA) plots, based on Bray-Curtis dissimilarity distances, show clustering patterns of the internal **(A)** and cuticle surface **(B)** microbiota in *An. albimanus* F_1_  larvae with respect to pyrethroid exposure. Separate plots, further sub-categorized by sample origin (depicted by unique shapes), are presented for each insecticide tested. Each point on the plots represents the bacterial composition of a pool of three mosquitoes, with the axes representing the first two dimensions of the PCoA, along with the proportion (%) of variation in bacterial composition captured. The plots show distinct separation of pyrethroid-exposed (resistant and susceptible) mosquitoes from -unexposed mosquitoes. Within the exposed group, the separation between resistant and susceptible mosquitoes varies from gradient to distinct. These clustering patterns are consistent for both pyrethroid insecticides tested and across all locations for the internal microbiota **(A)**. The cuticle surface microbiota **(B)** showed varying clustering patterns for each insecticide across locations. Results of overall non pairwise beta diversity (Bray Curtis) comparison using PERMANOVA (999 permutations) tests are presented. The test statistic value (pseudo-F) for each comparison is presented, with significance set to p-value <0.05. The difference in bacterial composition between resistant, susceptible and unexposed mosquitoes was not statistically significant in the internal microbiota **(A)**. Conversely, the bacterial composition on the cuticle surface was significantly different between all three categories for both deltamethrin and permethrin.

### *Asaia* dominated the internal and cuticle surface microbiota of adult *An. albimanus* but not larvae from the same population

Following taxonomic annotation of ASVs to the genus level, 75 and 140 bacterial genera were detected in adult and larvae, respectively; this was out of a total of 118 (adult) and 203 (larvae) assigned bacterial and archaeal taxa (Suppl. 9 and 10), with the archaeal reads only identified in adults. The less diverse adult microbiota predominantly comprised ASVs assigned to the genera *Asaia*, with >70% of these ASVs found in both the internal and cuticle surface microbial niches (Fig. 4). Conversely, ASVs assigned to *Asaia* comprised 0.02% of the overall larval microbiota, with *Leucobacter, Thorsellia*, and *Chryseobacterium* dominating the internal microbial niche (collectively constituting 35%), and *Acidovorax* and *Paucibacter* (each making up <50%) dominating the cuticle surface (Fig. 5 and Suppl. 10). Despite being comprised of >70% *Asaia,* the adult cuticle surface microbiota had nearly twice (n=106) as many taxa as the internal microbiota (n=62) (Suppl. 9). Out of the total adult microbial taxa detected (n=118), 47% were unique to the cuticle surface, 10% to the internal microbial niche, and 43% shared by both. In larvae, the cuticle surface microbiota was comprised of only slightly more taxa (n=194) compared to the internal (n=180) microbiota, and out of all taxa detected (n=203), 11% were unique to the cuticle surface, 4% to the internal microbial niche, and 85% were shared by both (Suppl. 10).

**Fig. 4.**
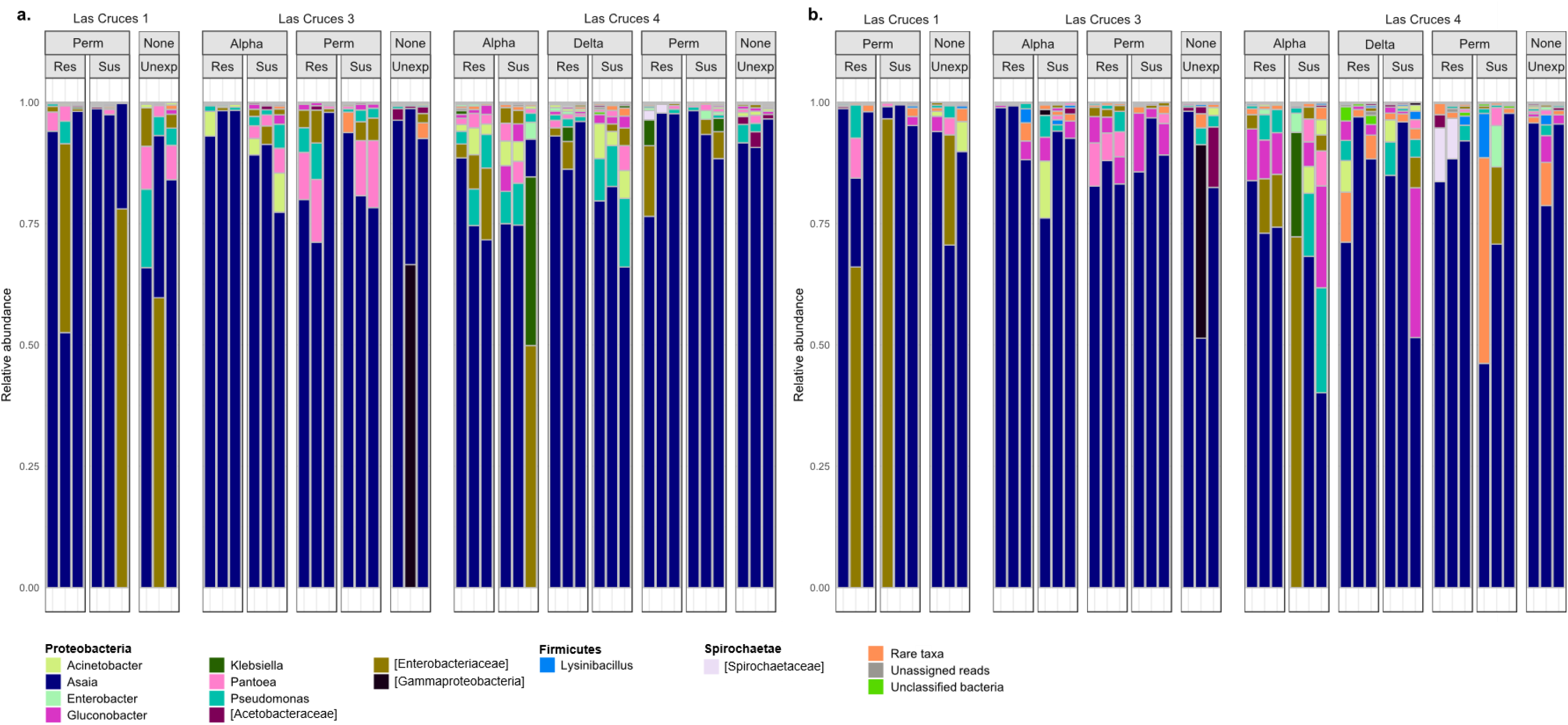
Barplots showing the relative abundance of taxonomically-annotated amplicon sequence variants from F_1_ adult *An. albimanus.* Amplicon sequence variances (ASVs) were taxonomically annotated to the genus level, and only taxa with relative abundance >0.1% are shown, all other taxa are collapsed and presented as ‘rare taxa’. The barplots show the relative abundance of annotated ASVs across all sites, sub-categorized by insecticide type and resistance status, indicating that the adult internal **(A)** and cuticle surface **(B)** microbiota is dominated by *Asaia*. Across all insecticides tested, the relative abundance of identified taxa differed between resistant, susceptible and unexposed mosquitoes in both internal **(A)** and cuticle surface **(B)** microbiota. ASVs that were not identified to the genus level are presented in square brackets, indicating the lowest possible level annotated. Identified taxa are organized by phylum, with phylum name indicated in bold. Res = resistant, Sus = susceptible and Unexp = non-exposed.

**Fig. 5.**
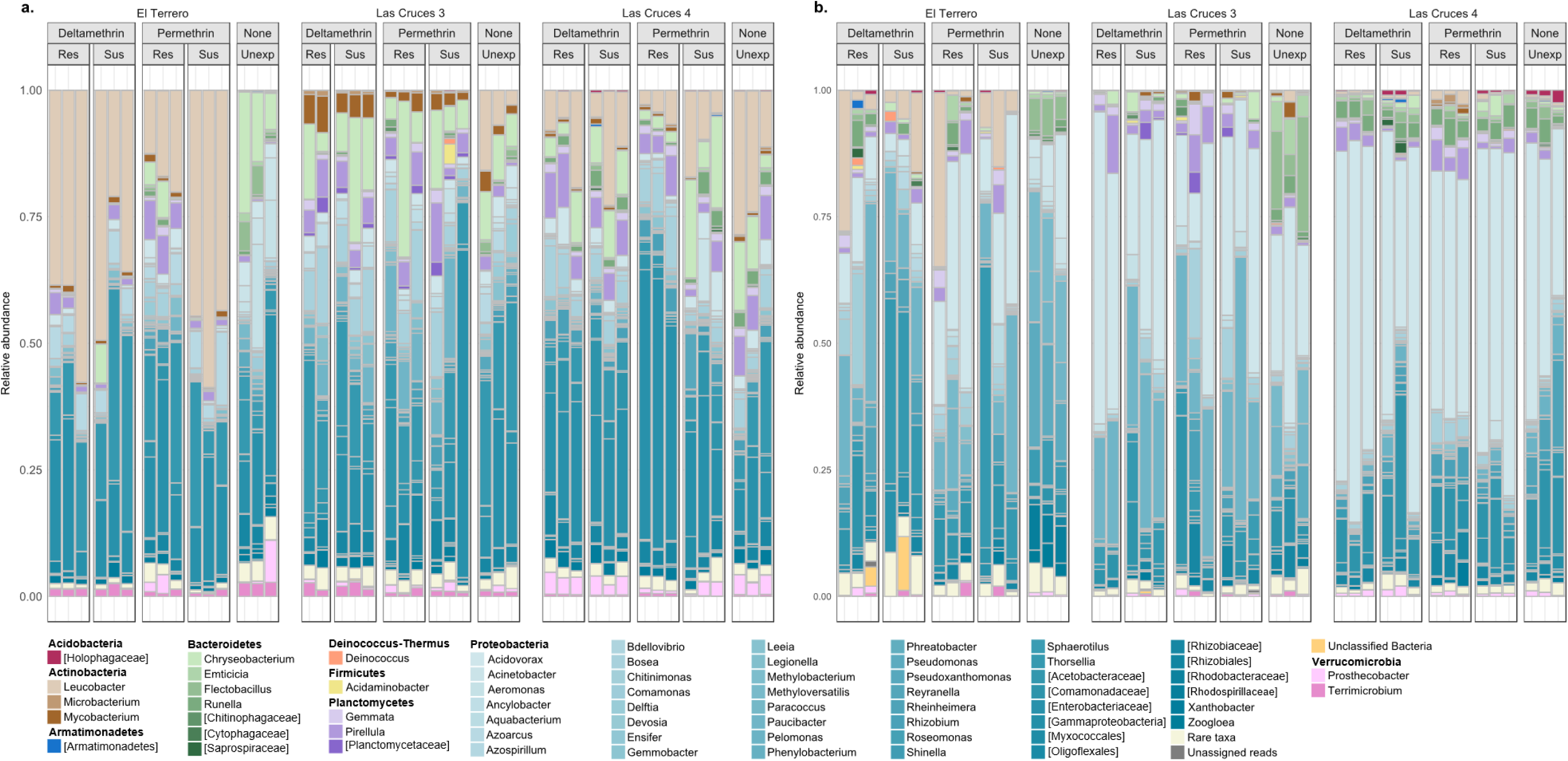
Barplots showing the relative abundance of taxonomically-annotated amplicon sequence variants from *An. albimanus* F_1_ larvae. Amplicon sequence variances (ASVs) were taxonomically annotated to the genus level, and only taxa with relative abundance >0.1% are shown, all other taxa are collapsed and presented as ‘rare taxa’. The barplots, grouped by location and sub-grouped by insecticide type and resistance status, show variable relative abundance of annotated ASVs across all locations, indicating that the larval internal **(A)** and cuticle surface **(B)** microbiota is mostly dominated by bacteria belonging to the phylum *Proteobacteria*. The genus *Leucobacter* was dominant in the internal microbiota, especially in El Terrero and Las Cruces 4 **(A)**, while *Acidovorax* dominated the cuticle surface microbiota **(B)**. Across all insecticides tested, the relative abundance of identified taxa differed between resistant, susceptible and non-exposed mosquitoes in both internal **(A)** and cuticle surface **(B)** microbiota. ASVs that were not identified to the genus level are presented in square brackets, indicating the lowest possible level annotated. Identified taxa are grouped and color-themed by phylum, with phylum name indicated in bold. Res = resistant, Sus = susceptible and Unexp = non-exposed

### *Klebsiella*, ***Asaia,* and *Pantoea* were consistently overabundant in pyrethroid resistant *An. albimanus***

The relative abundance of bacterial taxa in both larvae (Fig. 5) and adults (Fig. 4) differed between pyrethroid-resistant, -susceptible and non-exposed mosquitoes. This was consistent for each insecticide tested, and across all study sites. We consequently used the linear discriminant analysis (LDA) effect size (LEfSe) program (*30*) to identify bacterial genera that were consistently more abundant in resistant samples—these are subsequently referred to as possible bacterial markers of pyrethroid resistance.

*Klebsiella* was consistently over five times (*p*=0.02) more abundant in the internal microbiota of alphacypermethrin-resistant adults compared to -susceptible or non-exposed samples across the two locations tested (Fig. 6A). Similarly, *Asaia* (*p*=0.01) and *Pantoea* (*p*=0.03) were consistently over eight times more abundant in the internal microbiota of permethrin-resistant adults compared to -susceptible or non-exposed samples across all three locations tested (Fig. 6B). However, no bacterial genera in the internal microbiota of deltamethrin-resistant mosquitoes were more abundant than those of -susceptible or non-exposed mosquitoes. Rather, in deltamethrin-susceptible adults, *Chryseobacterium* was over five-fold more abundant when compared to resistant or non-exposed mosquitoes (Fig. 6C). Consistent with the results of the alpha diversity comparisons which showed that neither exposure to nor type of pyrethroid insecticide impacted bacterial abundance on the adult cuticle surface (Suppl. 8), no bacterial genus was significantly more abundant on the cuticle surface of resistant adult mosquitoes compared to susceptible or non-exposed mosquitoes, for any insecticide tested. Furthermore, for each individual insecticide tested, bacterial genera from neither the internal nor cuticle surface microbiota of resistant larvae were significantly more abundant than those of susceptible or non-exposed mosquitoes.

**Fig 6.**
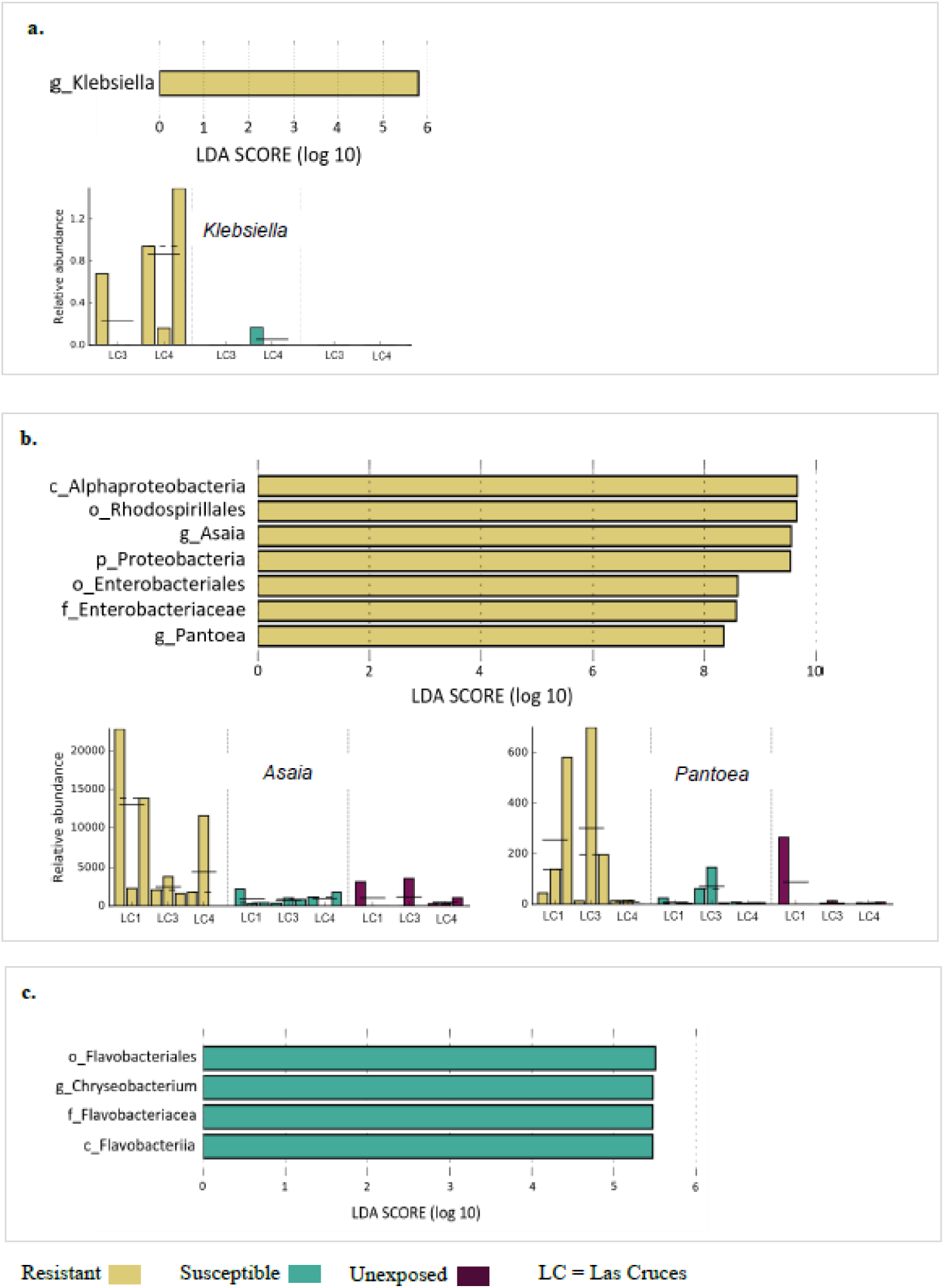
Linear discriminant analysis of possible bacterial markers of pyrethroid resistance in adult *An. albimanus*. The linear discriminant analysis effect size (LEfSe) tool was used to predict bacterial markers associated with pyrethroid resistance or susceptibility. This was based on identifying bacterial taxa that were consistently (across all sites tested) significantly more abundant (*p*<0.05 in pairwise Kruskal-Wallis and subsequent Wilcoxon rank-sum tests) in either resistant or susceptible mosquitoes compared to other categories. The horizontal histograms show fold change (log10 transformed) in relative abundance of bacterial taxa that were identified as possible markers, along with the relative abundance (vertical histograms) of each identified bacterial marker of pyrethroid resistance across all sites tested. The relative abundance of bacterial taxa was scaled using LEfSe’s 1,10^6^transformation prior to analysis. With over five fold more abundance in resistant compared to susceptible or non-exposed mosquitoes, *Klebsiella* was identified as a possible marker of alphacypermethrin resistance **(A)**, and *Asaia* and *Pantoea* were identified as possible markers of permethrin resistance **(B)** in adult internal microbiota. While no bacterial marker of deltamethrin resistance was identified, *Chryseobacterium* was over five fold more abundant in susceptible compared to resistant or non-exposed mosquitoes **(C)**.Taxonomic levels are indicated by p_ phylum; c_ class; o_ order; f_ family; g_ genus, with specific focus on the genus level. The mean and median relative abundance of each identified bacterial marker per location are indicated by solid and dashed horizontal lines on the relative abundance histograms respectively.

## Discussion

With increasing evidence of microbiota-mediated insecticide resistance in insects, particularly in agricultural pests (*13*), it is plausible that the mosquito microbiota could also contribute to the host’s insecticide detoxification processes. Building upon our earlier findings, which showed significant differences in bacterial composition between insecticide-resistant and –susceptible Peruvian *An. albimanus* (*10*), we characterized the microbiota of Guatemalan *An. albimanus* with differing pyrethroid insecticide resistance profiles. We focused specifically on late instar larvae (L3-L4) and adult (non-blood-fed virgin females) F_1_ progeny of field-caught mosquitoes to obtain uniform physiological profiles, while maintaining the genetic background of the field populations from where they were collected. Our results showed that insecticide exposure, insecticide type, microbial niche (internal or cuticle surface microbiota) and host developmental stage significantly impact *An. albimanus* microbiota. We present the first description, to our knowledge, of the effects of pyrethroid insecticides on mosquito microbiota, and the first description, of the microbiota of *An. albimanus* larvae. We also characterized the microbiota on the mosquito cuticle surface for the first time.

Overall bacterial composition differed significantly between pyrethroid-exposed and non-exposed larvae and adults. Considering that the cuticle surface is the first site of insecticide contact, we had anticipated a general effect of insecticide exposure, if present, on the cuticle surface microbiota of both larvae and adults. While this was indeed what we observed in larvae, the result was the opposite for adult mosquitoes, with an effect only detected on the internal microbiota. Our results also showed significant differences in bacterial abundance and evenness driven by insecticide type and insecticide exposure, in adult internal, but not cuticle surface microbiota. This insecticide type-and exposure-driven difference in bacterial abundance and evenness was also evident in larvae, but with respect to the cuticle surface microbiota rather than internal microbiota. The differing impact of insecticide type and exposure on larval and adult microbiota could be explained by the differences in how mosquito larvae and adults come into contact with insecticides. Throughout the bioassays, larvae were fully immersed in insecticide-treated water, mimicking their natural exposure to insecticides. Adults were introduced into insecticide-coated bottles (also mimicking natural insecticide exposure) where they mainly contacted the insecticide via their tarsi. As such, the larger insecticide-contact area (the entire body surface), as well as their prolonged contact with insecticide, could have led to the impact on the composition of the cuticle surface microbiota in larvae. Adult malaria vectors typically pick up insecticides on their tarsi upon landing on treated surfaces (*33*), which may explain the insignificant effect of insecticide exposure on their cuticle surface microbiota. It is not immediately apparent how pyrethroid insecticide exposure impacts the internal microbiota of adult mosquitoes. In insects, pyrethroid insecticides are often metabolized by mixed function oxidase enzymes (*34*), a characteristic component of internal mosquito tissues including the midgut (*35*), where they are preferentially produced (*36*). The midgut is also where the majority of the internal microbiota is found, so the microbes within this tissue (along with other internal tissues) could conceivably be impacted by exposure to pyrethroid insecticides. The internal microbiota of insecticide-exposed adult mosquitoes could also reflect the presence of pre-selected microbiota that is linked to the insecticide resistance status of the host. When individual pyrethroid insecticides were considered, the effects of insecticide exposure were generally consistent across insecticides except for deltamethrin, where no significant effect of exposure was detected on either internal or cuticle surface bacteria in both larvae and adults. Furthermore, in alphacypermethrin-exposed adults, insecticide exposure significantly impacted both internal and cuticle surface bacterial composition. The impact of exposure on the cuticle surface microbiota of alphacypermethrin-exposed adults could indicate a more intense resistance to alphacypermethrin in this mosquito population compared to other insecticides. Indeed, bioassays indicated a higher proportion of alphacypermethrin resistance in this mosquito population compared to other insecticides (Suppl. 1C).

Congruent with the effects of insecticide type and insecticide exposure on the mosquito microbiota, we identified consistently overabundant bacterial genera in the internal microbiota of resistant adult mosquitoes. These were *Klebsiella, Pantoea* and *Asia*, and are considered as possible bacterial markers of pyrethroid resistance in adult *An. albimanus*. Consistent with results from our previous study (*10*), these bacterial genera were more abundant in the internal microbiota of insecticide-resistant compared to -susceptible or non-exposed adult mosquitoes. In alphacypermethrin-exposed adults, across the two locations tested, *Klebsiella* was over five-fold more abundant in resistant compared to susceptible or non-exposed samples. It is well-established that bacteria belonging to the genera *Klebsiella* metabolize insecticides (*37, 38*), including pyrethroids (*9*). In fact, a pyrethroid hydrolyzing esterase, EstP, has been identified in *Klebsiella*, and its encoding gene has been characterized (*9*). Nonetheless, the role of *Klebsiella* in pyrethroid metabolism in insect hosts remains to be functionally validated. In the internal microbiota of permethrin-resistant adults, *Pantoea* and *Asaia* were over eight-fold more abundant in contrast to the microbiota of susceptible or non-exposed mosquitoes. Unlike *Klebsiella*, limited information is available about the role of *Pantoea* in metabolizing insecticides within insects, but our previous identification of a higher proportion of this bacterial genera in insecticide resistant (compared to susceptible) mosquitoes suggests that they could potentially be contributing to insecticide resistance (*10*). Similarly little has been documented about insecticide metabolism in *Asaia*, but pyrethroid-metabolizing enzymes have been shown to be preferentially produced in several primary detoxification tissues in mosquitoes, including the midgut (*35*), where the majority of *Asaia* are known to reside (*39*). One possibility is that when insecticides penetrate the midgut (and other internal tissues) they could be metabolized by the resident microbes, including *Asaia*. As a paratransgenesis candidate (*39*), *Asaia*’s potential role in insecticide resistance should not be overlooked. In contrast however, no possible bacterial markers were identified in either internal or cuticle surface larval microbiota, and a transient larval microbiota (*40, 41*) could be one reason for this.

Insect internal microbiota have been shown to contribute to the metabolism of topically applied insecticides, consequently contributing to host insecticide resistance (*13*). Our results show a significant impact of pyrethroid exposure on the bacterial composition of mosquito microbiota, and we have also identified significantly more abundant pyrethroid-metabolizing bacteria in resistant adults, consistently across multiple locations. This indicates several events that may be occurring within the host, perhaps simultaneously; whereby insecticide exposure influences the mosquito microbiota by selecting for insecticide-metabolizing bacteria, and these bacteria contribute to insecticide detoxification within the host, consequently contributing to the host’s overall resistance to insecticides. The results presented here, using adult F_1_ progeny from field-caught mosquitoes, along with those from our previous work on field-caught adult mosquitoes (*10*), suggest that insecticide exposure could be selecting for insecticide-metabolizing bacteria. This may be particularly important with respect to insecticide resistance intensity. Our previous findings showed significant differences in bacterial composition between mosquitoes that were susceptible to the diagnostic dose of fenitrothion and those that were resistant to five times the diagnostic dose (*10*). However, in the present study, we only considered mosquitoes that were susceptible or resistant to the diagnostic dose of the insecticides due to low insecticide resistance intensity in the study area. Regardless, exposure to insecticides significantly impacted bacterial composition, indicating that ongoing insecticide exposure could continue to select for insecticide-metabolizing bacteria. This could be of even greater relevance in areas where the intensity of resistance is high. The underlying mechanisms of insecticide resistance intensity in mosquitoes are poorly understood, and we propose that the metabolism of insecticides by host microbiota could be a contributing factor.

While no bacterial markers of deltamethrin resistance were identified in the internal microbiota of adult mosquitoes, *Chryseobacterium*—previously named *Flavobacterium*—was over five-fold more abundant in susceptible, compared to resistant or non-exposed samples. Bacteria belonging to the genus *Chryseobacterium* (*Flavobacterium*) have previously been identified in *Anopheles* mosquitoes, including lab-reared *An. albimanus* (*42*), however their role in mosquito physiology has not been described.

As has previously been shown (*31*), larval microbiota was more diverse than adult microbiota in this study, and overall, the identified bacterial taxa have previously been identified in *Anopheles* mosquitoes (*12, 43*), including Latin American *Anopheles* (*10, 43*). While we characterized the bacterial composition of *An. albimanus* larvae for the first time in this study, our results included bacterial taxa that have been identified in other *Anopheles spp.* larvae (*31, 44*). Unlike in humans where the skin surface microbiota is well characterized, studies on insect cuticle surface is sparse. Human skin surface comprise less diverse bacterial communities compared to the internal organs (collectively) (*45*). Similarly, a survey of the cuticle surface microbiota of Canadian dark beetles revealed a more diverse internal microbiota compared to the cuticle surface (*46*). However, in this study, we report that in both mosquito larvae and adults, the cuticle surface contains more diverse bacteria compared to the internal microbial niche. Lower diversity of internal microbiota could indicate a conserved internal bacterial community in mosquitoes, as have previously been reported, thus corroborating earlier reports of the presence of an internally selective environment (*31*). While our data showed that the mosquito cuticle surface harbors a more diverse bacterial community compared to the internal microbial niche, 85% and 43% of the bacterial taxa were shared by both internal and cuticle surface microbiota in larvae and adults, respectively. This suggests that bacteria on the cuticle surface are not only incidentally acquired from the host’s environment. The relationship between bacterial composition of the mosquito cuticle surface and the host environment remains poorly understood, and it remains unknown how the cuticle surface microbiota may contribute to the host’s physiological processes.

We also report the detection of *Asaia* in *An. albimanus* for the first time, and showed that it dominated (>70%) the adult internal and cuticle surface microbiota. While laboratory raised *Anopheles* mosquitoes have been shown to predominantly harbor *Asaia* (*39*), studies identifying *Asaia* in field-collected *Anopheles* mosquitoes have commonly been qualitative (*47, 48*), with sparse documentation of variable *Asaia* abundance. *Asaia* was also present in larval microbiota, albeit in negligible proportions (0.02%), (noting that the larvae tested were from the same parents and generation as the adults tested). Thus, supporting the notion that while the microbiota may be transient in immature mosquito stages due to multiple molting events, rapid development and physiological changes (*41*), some bacteria can be transstadially transmitted to the adult stage (*7*). Furthermore, the predominance of *Asaia* in adults despite their negligible proportions in larvae is strongly indicative of *Asaia*’s ability to quickly and efficiently colonize and dominate the microbiota of adult *Anopheles* mosquitoes (*39*), thus resulting in its consideration as a paratransgenesis candidate for malaria control (*49*). Moreover, since we used non-blood fed virgin adult females in this study, the overabundance of *Asaia* in adults (which may also have been acquired independently of the larval stage) could be an indicator of the host’s age and/or feeding status. This is because mosquito age and feeding status are known to impact bacterial composition (*50*), and *Asaia* is a known component of sugar sources (*51*). Our sequencing approach deeply sampled the mosquito microbiota, as indicated by the plateaued rarefaction curves (Suppl. 4), thus identifying bacterial taxa in each sample beyond the point at which further sequencing had no impact on the abundance of detected bacterial communities. The disparity detected in abundance of *Asaia* between larvae and adults is thus biologically relevant and not a sequencing artifact. While a few studies have shown that the mosquito microbiota varies across host’s developmental stages (*31, 40*), more work is still needed to understand when and how the mosquito microbiota changes throughout host development, and how these changes might affect host physiology with regard to insecticide resistance and other key characteristics, such as vector competence.

The results presented here highlight differential effects of insecticide exposure on the mosquito microbiota across mosquito developmental stages and insecticide types, and indicate the presence of a conserved microbiota—particularly within adult mosquitoes—that is altered by insecticide exposure. These results also identify overabundant insecticide-metabolizing bacteria in pyrethroid resistant mosquitoes that could be possible markers of resistance in field populations of *An. albimanus*. Together these findings indicate insecticide selection pressures on mosquito microbiota, and support the hypothesis of a microbe-mediated mechanism of insecticide metabolism in mosquitoes. Focusing on the potential bacterial markers identified in the current study, future work will characterize specific bacterial components that are being affected by pyrethroid exposure and/or contributing to resistance in mosquito populations.

## Acknowledgments

This work is supported by the US Centers for Disease Control and Prevention (CDC) through the American Society for Microbiology’s (ASM) Infectious Disease and Public Health Microbiology Postdoctoral Fellowship program, and the CDC’s Advanced Molecular Detection (AMD) program. We thank the Malaria Research and Reference Reagent Recourse Center (MR4) for providing the ITS2 primers used for *Anopheles albimanus* species identification; Nelson Jimenez, Ricardo Valle and Ricardo Santos from the Ministerio de Salud Publica y Asistencia Social (MSPAS) for assistance during mosquito sampling; Daniela Da’Costa, Pedro Peralta, Adel Mejia and Alfonso Salam from Universidad del Valle de Guatemala (UVG) for field support and assistance during mosquito rearing and insecticide resistance assays; Stephen Smith from the Entomology Branch, CDC, for insightful discussions on pyrethroid chemistry and mosquito cuticle permeability; and Numi T for the fortitude offered throughout the data analysis and preparation of the manuscript.

## List of supplementary Materials

Suppl. 1. Insecticide bioassay data

Suppl. 2. Sequencing output statistics

Suppl. 3. Variables included in regression model

Suppl. 4. Shannon diversity rarefaction curves

Suppl. 5. Alpha diversity box plots showing effects of insecticide type and exposure on bacterial abundance and evenness

Suppl. 6. Beta diversity comparison of bacterial composition between pyrethroid-exposed and non-exposed *An. albimanus*

Suppl. 7. Pairwise beta diversity comparison showing effects of exposure to each insecticide on bacterial composition

Suppl. 8. Pairwise beta diversity comparisons showing effects of insecticide type on bacterial composition

Suppl. 9. Adults ASV Counts and relative abundance of bacterial taxa

Suppl. 10. Larvae ASV Counts and relative abundance of bacterial taxa

