## Supplementary material for "Pyrethroid exposure alters *Anopheles albimanus* microbiota and resistant mosquitoes harbor more insecticide-metabolizing bacteria": Suppl. 2

**Suppl. 2: Sequencing outputs and proportion of reads used for downstream analysis following quality control and dereplication.**

| | | | Number of reads (% remaining following QC, dereplication and chimera removal) | | | | | |
|---|---|---|---|---|---|---|---|---|
| | | | Internal microbiota | | | Cuticle surface microbiota | | |
| **Stage** | **Location** | **Insecticide** | **Resistant** | **Susceptible** | **Unexposed** | **Resistant** | **Susceptible** | **Unexposed** |
| **Adults** | Las Cruces 1 | Permethrin | 2495656 (60) | 2421158 (59) | 2356223 (61) | 1847318 (64) | 1973328 (58) | 1220490 (75) |
| | Las Cruces 3 | Alphacypermethrin | 2575153 (75) | 2279853 (66) | 2697153 (71) | 1451582 (85) | 1602746 (80) | 1687593 (85) |
| | | Permethrin | 2434964 (62) | 2035362 (62) | | 1685611 (85) | 1723938 (89) | |
| | Las Cruces 4 | Alphacypermethrin | 2403001 (65) | 2224020 (48) | 1996979 (79) | 1548342 (81) | 1905025 (45) | 1379612 (91) |
| | | Deltamethrin | 1657741 (74) | 1999803 (66) | | 1018949 (85) | 1208000 (82) | |
| | | Permethrin | 1552101 (62) | 1030681 (69) | | 1437977 (87) | 1350102 (79) | |
| **Total** | | | **13118616 (66)** | **11990877 (61)** | **7050355 (70)** | **8989779 (80)** | **9763139 (70)** | **4287695 (84)** |
| **Larvae** | El Terrero | Deltamethrin | 2285602 (36) | 2417346 (34) | 3160044 (26) | 807512 (46) | 174347 (34) | 2042499 (35) |
| | | Permethrin | 2440043 (31) | 2468291 (35) | | 1732541 (50) | 1054643 (47) | |
| | Las Cruces 3 | Deltamethrin[a] | 1187935 (35) | 2698429 (30) | 2983923 (30) | 1203630 (57) | 1671137 (48) | 2740549 (44) |
| | | Permethrin | 2266161 (33) | 2651416 (32) | | 1656194 (45) | 1629694 (47) | |
| | Las Cruces 4 | Deltamethrin | 1387104 (35) | 3239030 (25) | 2768185 (30) | 2005520 (66) | 1589326 (60) | 1505523 (55) |
| | | Permethrin | 2120366 (33) | 2636276 (33) | | 1760717 (57) | 1731822 (61) | |
| **Total** | | | **11687211 (34)** | **16110788 (31)** | **8912152 (28)** | **9166114 (54)** | **7850969 (53)** | **6288571 (44)** |

[a] Only two pools of larvae were processed for the deltamethrin resistant category

We characterized the internal and cuticle surface microbiota of whole larvae (n=132) and adult (135) $F_1$ progeny of wild caught *Anopheles albimanus* from four locations in La Gomera, Guatemala using 16S rRNA gene sequencing. These mosquitoes had been categorized as resistant, susceptible or unexposed to pyrethroid insecticides using the CDC bioassay, and for sequencing, were consolidated into 44 and 45 pools of larvae and adults respectively. Each pool comprised 3 mosquitoes, and 3 pools were processed per category (i.e. resistant, susceptible or unexposed). A total of 55,200,461 (adults) and 60,015,805 (larvae) sequencing reads were obtained and processed for downstream analysis. The table provides a breakdown of sequencing reads per category and the proportion of reads used for downstream analysis.
