## Supplementary material for "Pyrethroid exposure alters *Anopheles albimanus* microbiota and resistant mosquitoes harbor more insecticide-metabolizing bacteria": Suppl. 3

**Suppl. 3: variables included in regression model**

| S/N | Variables | Description | Categories |
|---|---|---|---|
| 1 | Location | Collection site of parent population | Las Cruces 1, Las Cruces 3, Las Cruces 4 or El Terrero |
| 2 | Phenotype | Pyrethroid resistance phenotype based on outcomes of | Resistant, Susceptible, or Unexposed |
| 3 | Type of insecticide | Type of pyrethroid insecticide tested | permethrin, alphacypermethrin or deltamethrin |
| 4 | Insecticide exposure | | Exposed (i.e. Resistant & Susceptible) or unexposed |
| 5 | Developmental stage | | Larva (L3-L4) or adult (2-5 d) |
| 6 | Microbial Niche | Collection site of the microbiota | Cuticle surface or internal |
