## Supplementary material for "Pyrethroid exposure alters *Anopheles albimanus* microbiota and resistant mosquitoes harbor more insecticide-metabolizing bacteria": Suppl. 4

### Suppl. 4: Rarefaction curves & depth

The following figures show rarefaction curves of Shannon alpha diversity index for each mosquito pool (3 mosquitoes/pool) processed by developmental stage, microbial niche and type of insecticide tested. All curves plateaued, indicating that additional sampling efforts did not result in changes in abundance and evenness of microbial taxa per sample. At each sampling depth shown, each curve shows the average Shannon diversity value, along with the range (boxplots—minimum, median and maximum) of values from 10 rarefaction iterations

### Suppl. 4A. Alphacypermethrin: adult cuticle surface

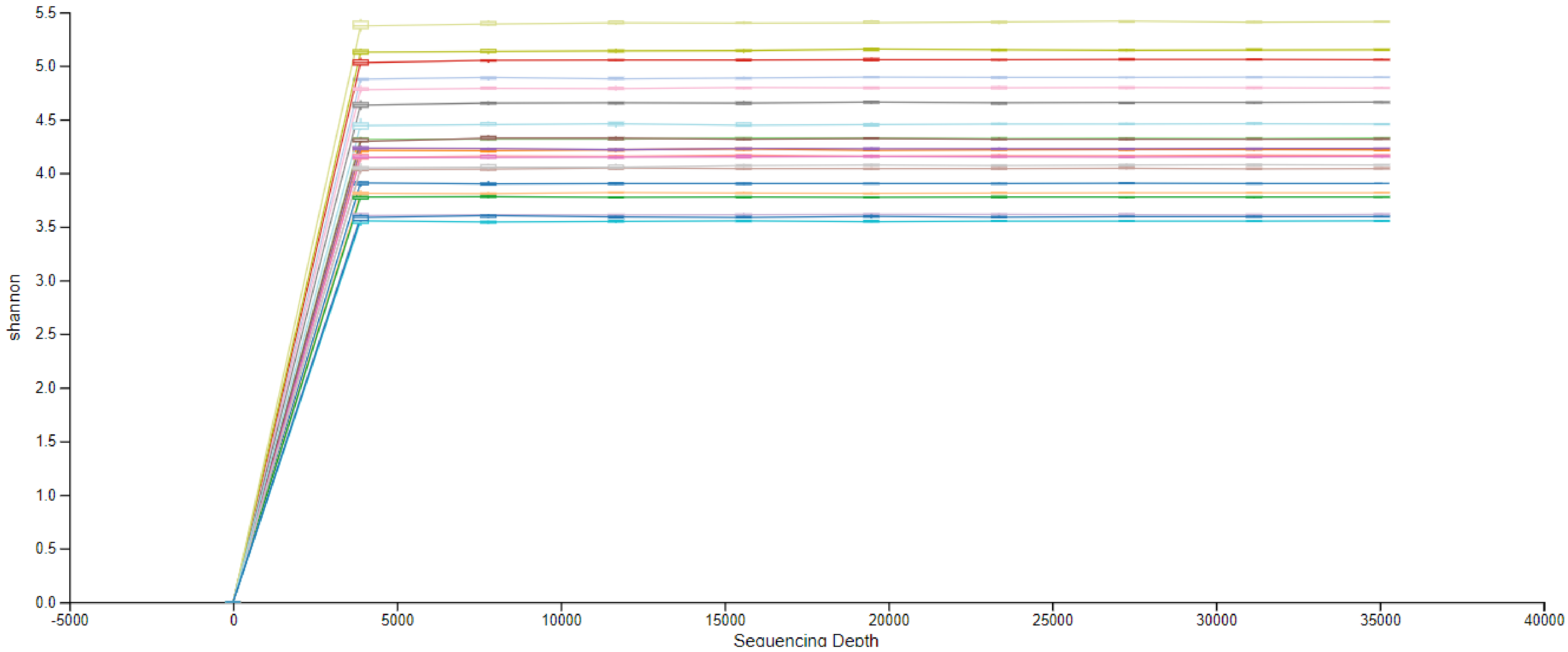

### Suppl. 4B. Alphacypermethrin: adult internal microbiota

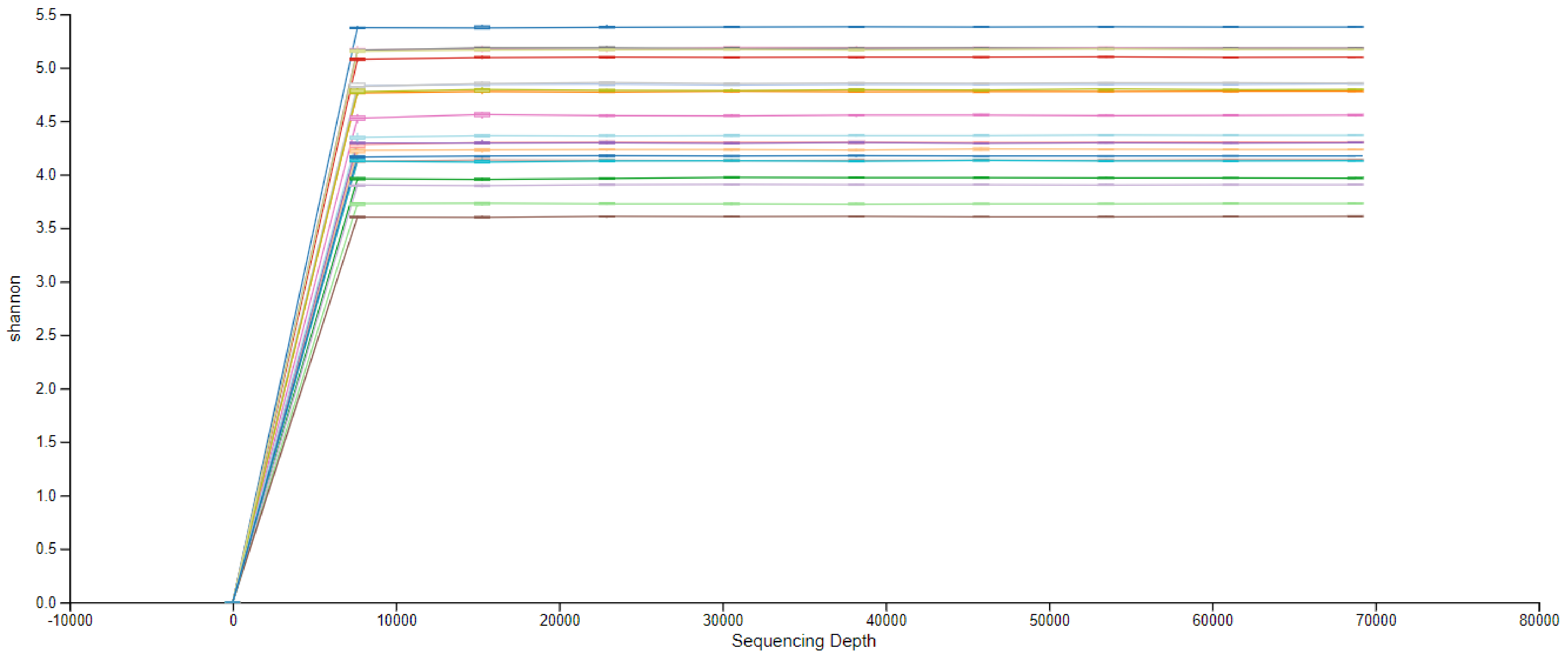

### Suppl. 4C. Deltamethrin: adult cuticle surface

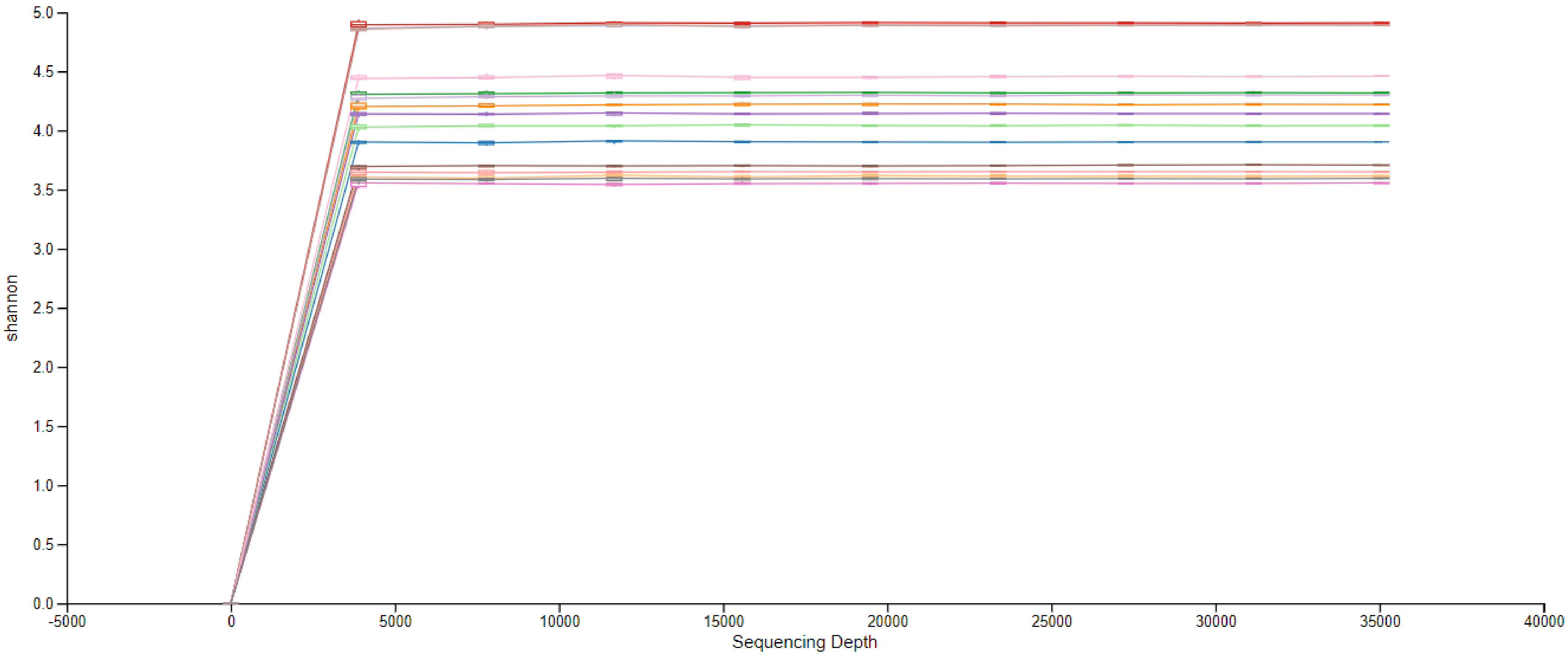

### Suppl. 4D. Deltamethrin: adult internal microbiota

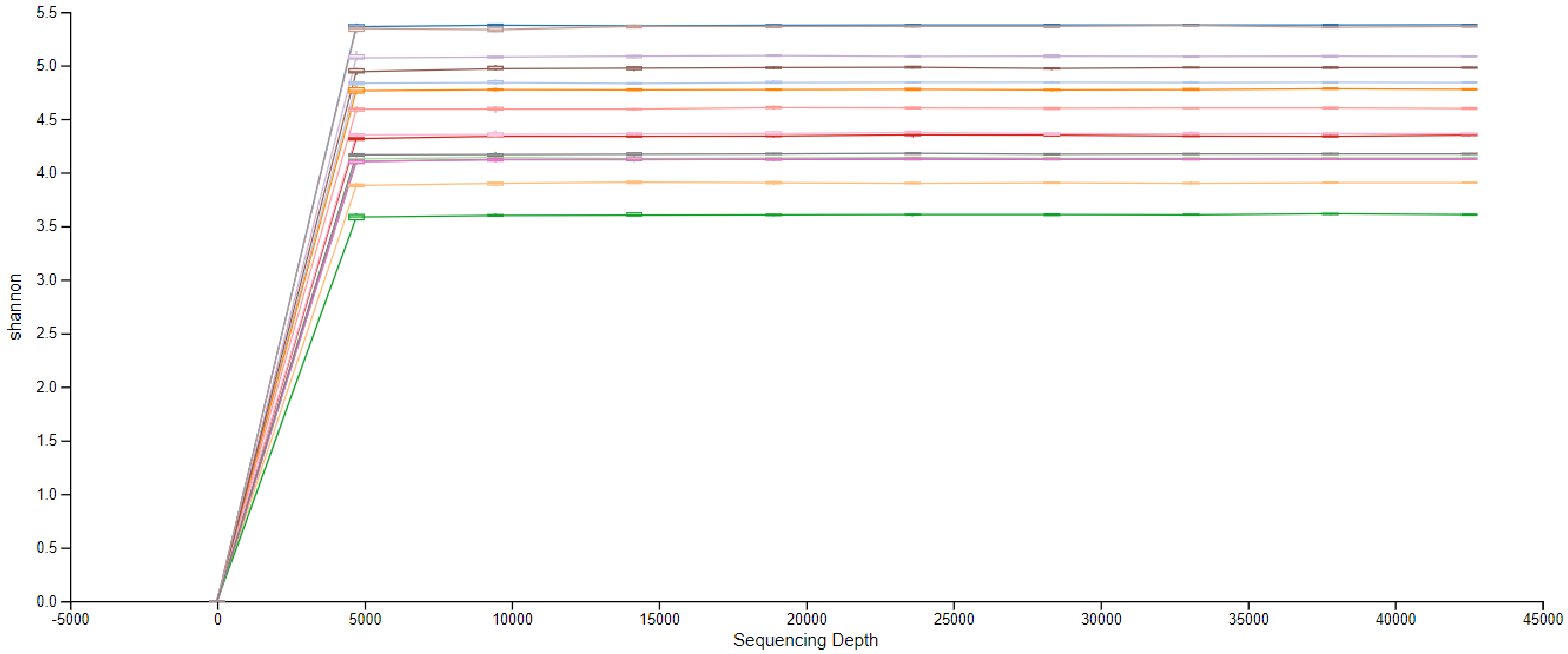

### Suppl. 4E. Permethrin: adult cuticle surface

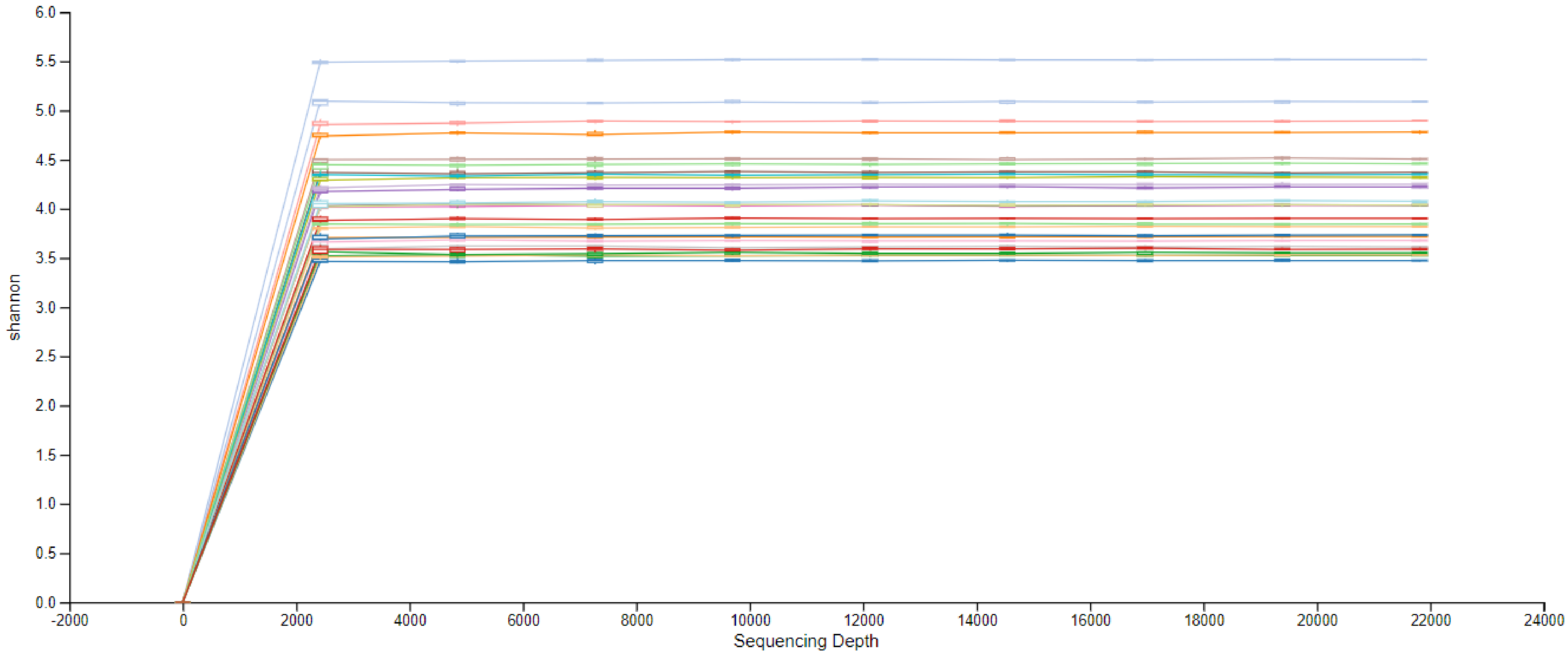

### Suppl. 4F. Permethrin: adult internal microbiota

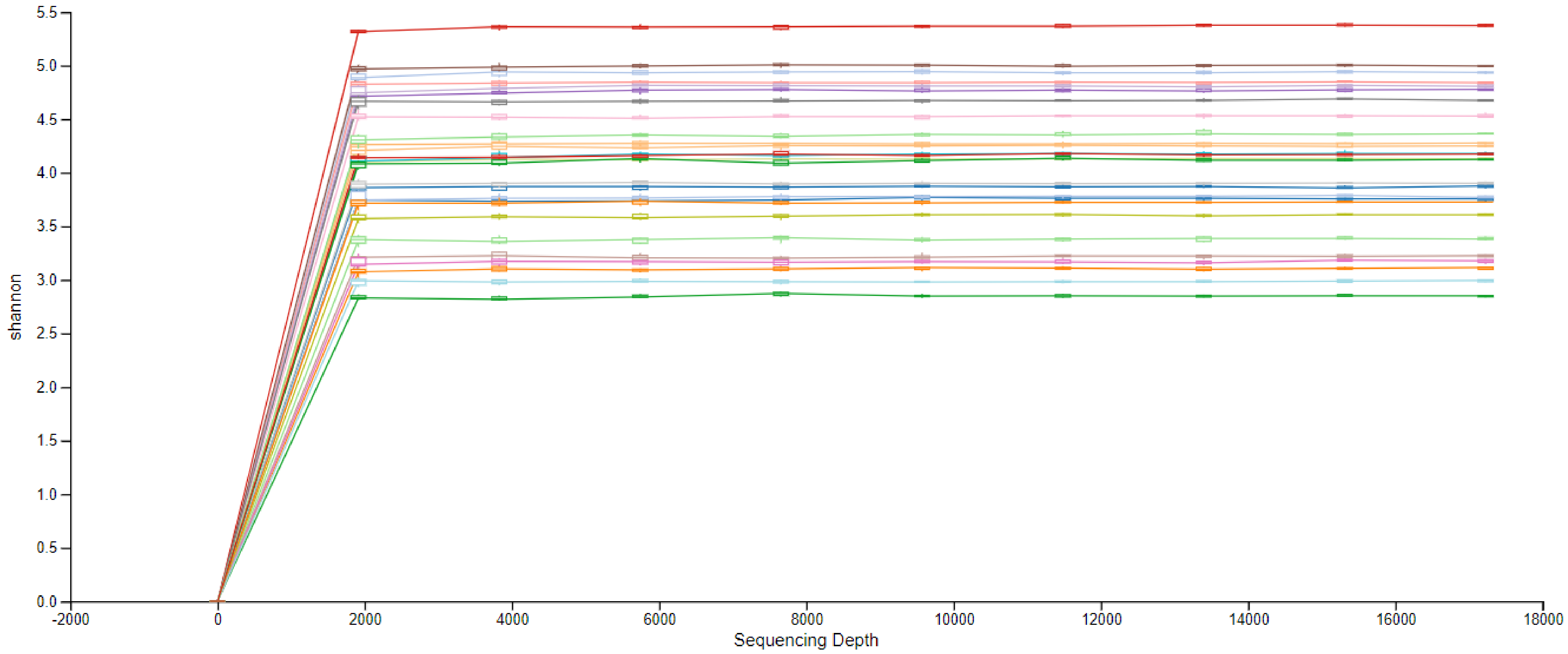

### Suppl. 4G. Deltamethrin: larval cuticle surface

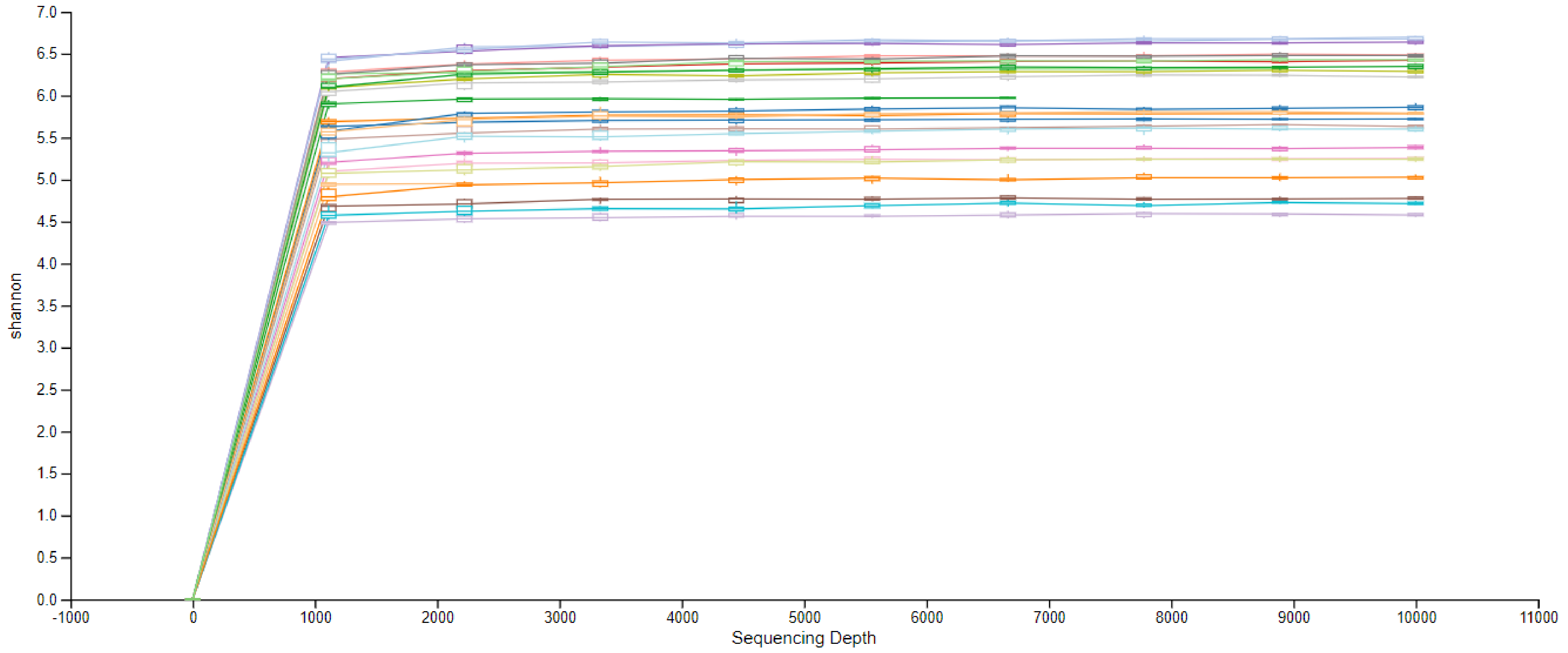

### Suppl. 4H. Deltamethrin: larval internal microbiota

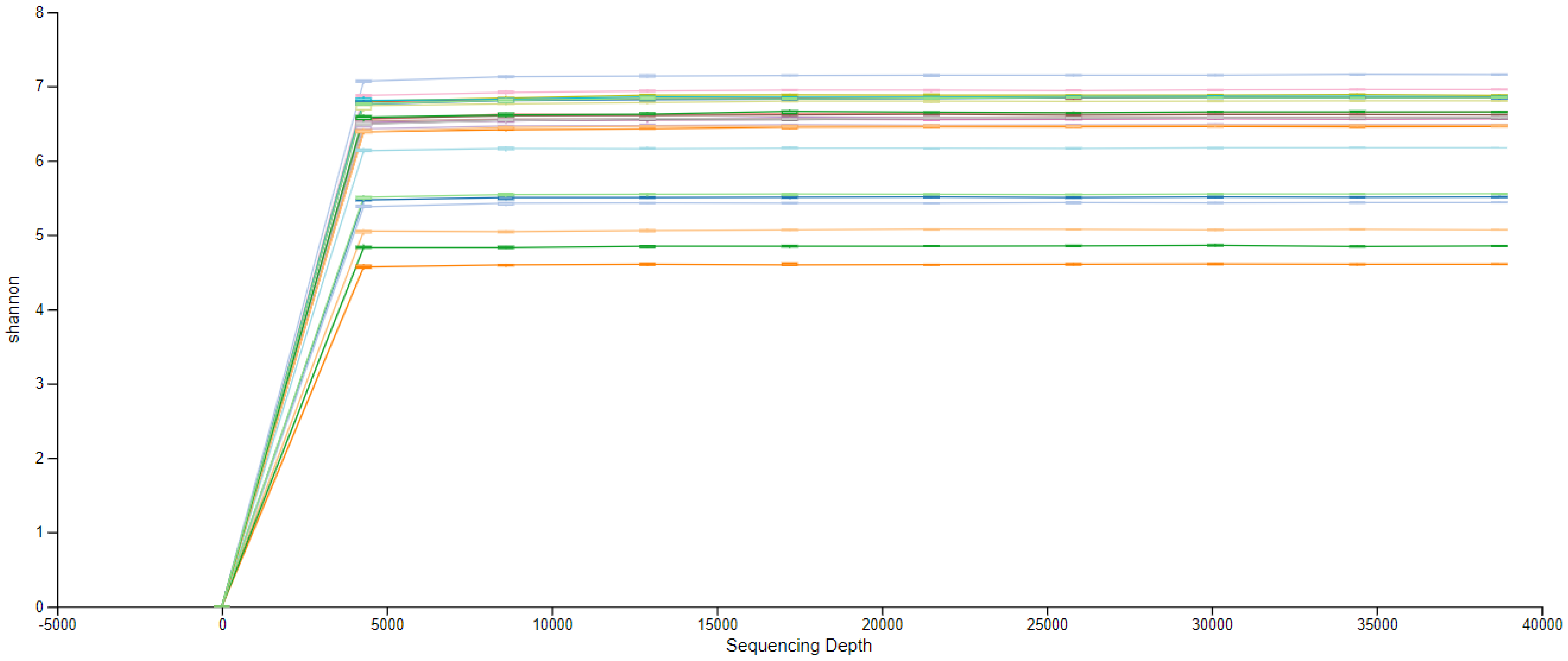

### Suppl. 4I. Permethrin: larval cuticle surface

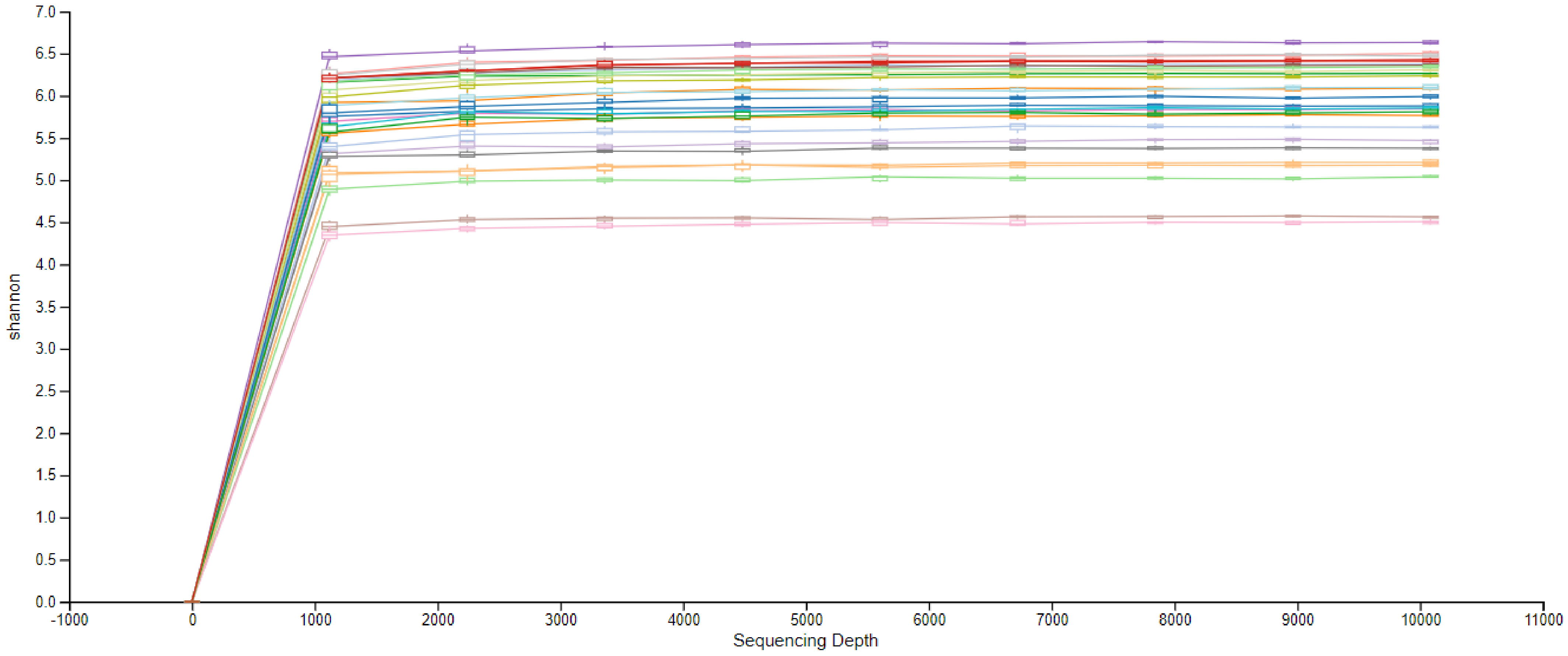

### Suppl. 4J. Permethrin: larval internal microbiota

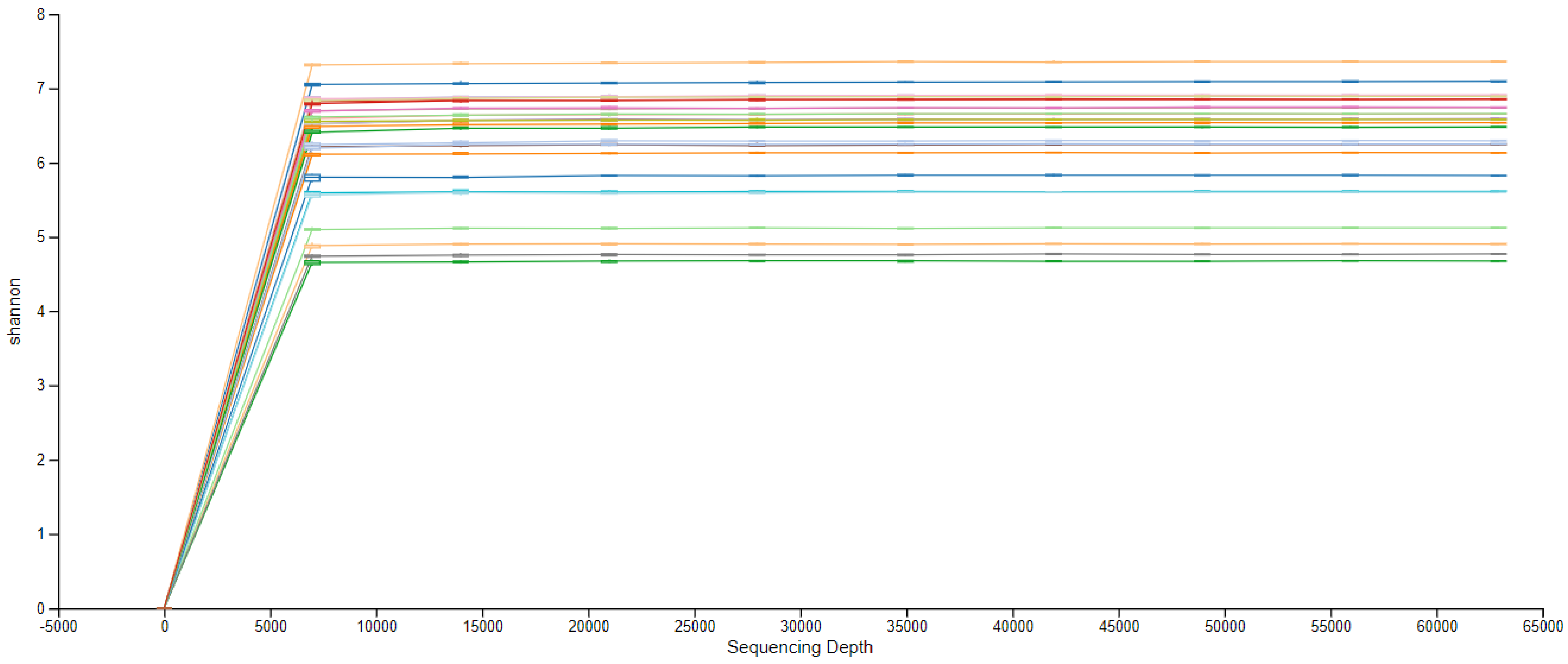

Suppl. 4k. Depth of rarefaction across categories.

Sequencing reads ranged from 3,277 - 223,222 across samples, and to obtain an even sampling depth for alpha diversity analysis, samples were rarefied to at least 10000 reads/sample across categories - a depth that was sufficient to capture the overall  composition of each sample. In categories with more reads, rarefaction was set to the lowest number of reads in that category.

| | | Rarefaction depth | |
|---|---|---|---|
| Stage | Insecticide | Internal microbes | Microbes on surface of the cuticle |
| Adults | Alphacypermethrin | 68781 | 35066 |
| | Deltamethrin | 42524 | 35066 |
| | Permethrin | 17227 | 21817 |
| Larvae | Deltamethrin | 38715 | 10000* |
| | Permethrin | 62928 | 10086 |

*lowest number of reads in this categpry was 3277. Three pools of delatamethrin susceptible samples from El Terrero were excluded from diversity analysis at this depth
