## Supplementary material for "Pyrethroid exposure alters *Anopheles albimanus* microbiota and resistant mosquitoes harbor more insecticide-metabolizing bacteria": Suppl. 5

### Suppl. 5. Alpha diversity comparisons

Boxplots showing alpha-diversity comparisons performed using the Shannon diversity indices of adult and larvae *Anopheles albimanus* microbiota. Whiskers on boxplots represent the range of minimum and maximum Shannon index values within each group, and grey dots represent outliers. Comparisons were performed using Kruskal-Wallis pair-wise tests (H) with Benjamini–Hochberg FDR correction (q-value). Significance was set to $q<0.05$

n, number of mosquito pools (3 mosquitoes/pool)

**Suppl. 5A. Bacterial abundance and evenness on the cuticle surface of *An. albimanus* larvae** was significantly impacted by exposure to either deltamethrin or permethrin insecticide, H=11.9, p=0.003. Larvae that were not exposed to insecticides represented the most abundant and even bacterial community, and no significant difference in bacterial abundance and evenness was seen between deltamethrin and alphacypermethrin tested mosquitoes

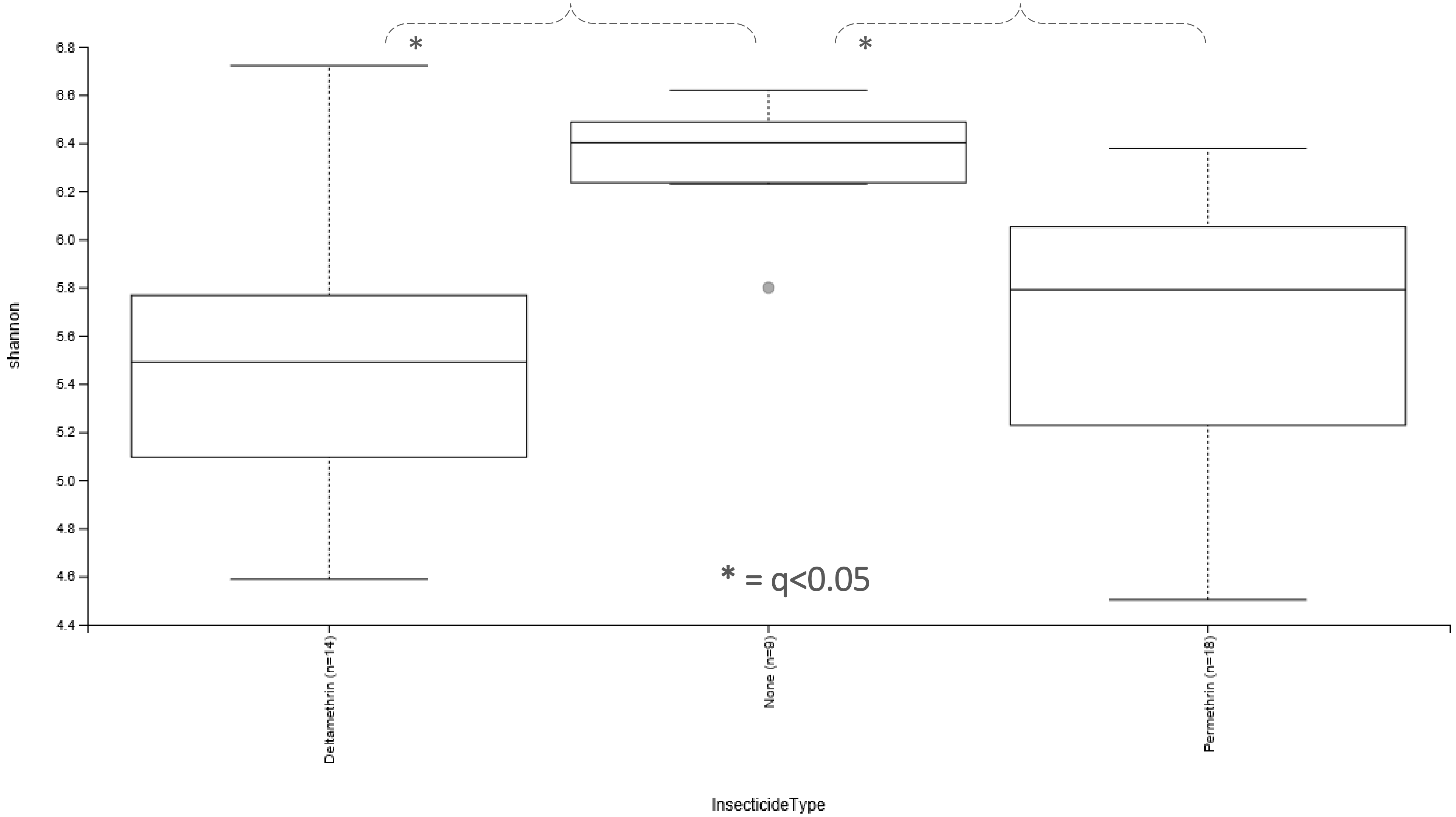

Suppl. 5B. Type of insecticide did not impact internal bacterial abundance and evenness in *An. albimanus* larvae. H = 2.7, p=0.25.

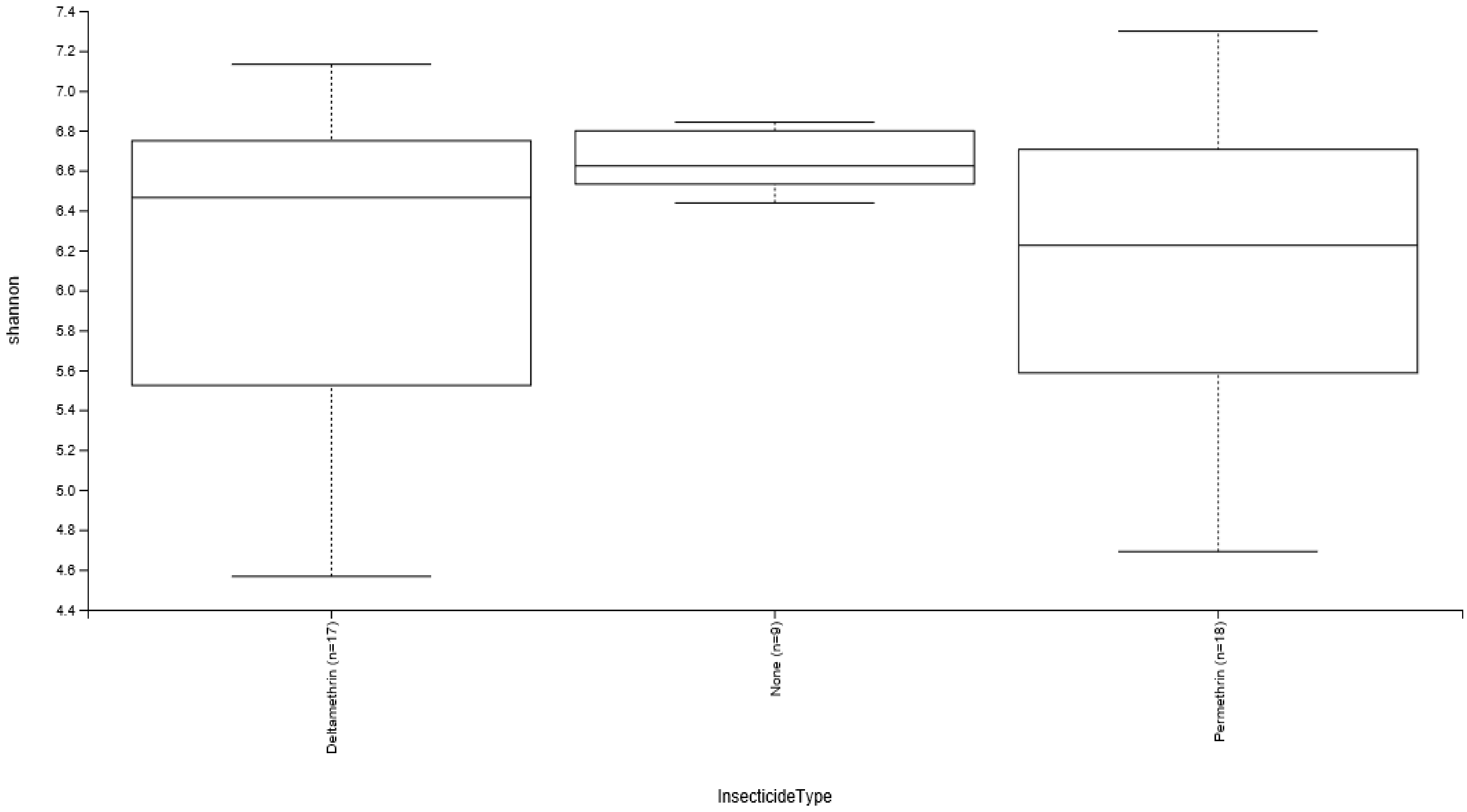

**Suppl. 5C. Insecticide exposure significantly impacts the abundance and evenness of bacteria on the cuticle surface of *An. albimanus* larvae**, H=12.3, p=0.002. Larvae that were not exposed to insecticides represented the most abundant and even bacterial community, with bacterial abundance and evenness differing significantly between unexposed and either resistant or susceptible mosquitoes

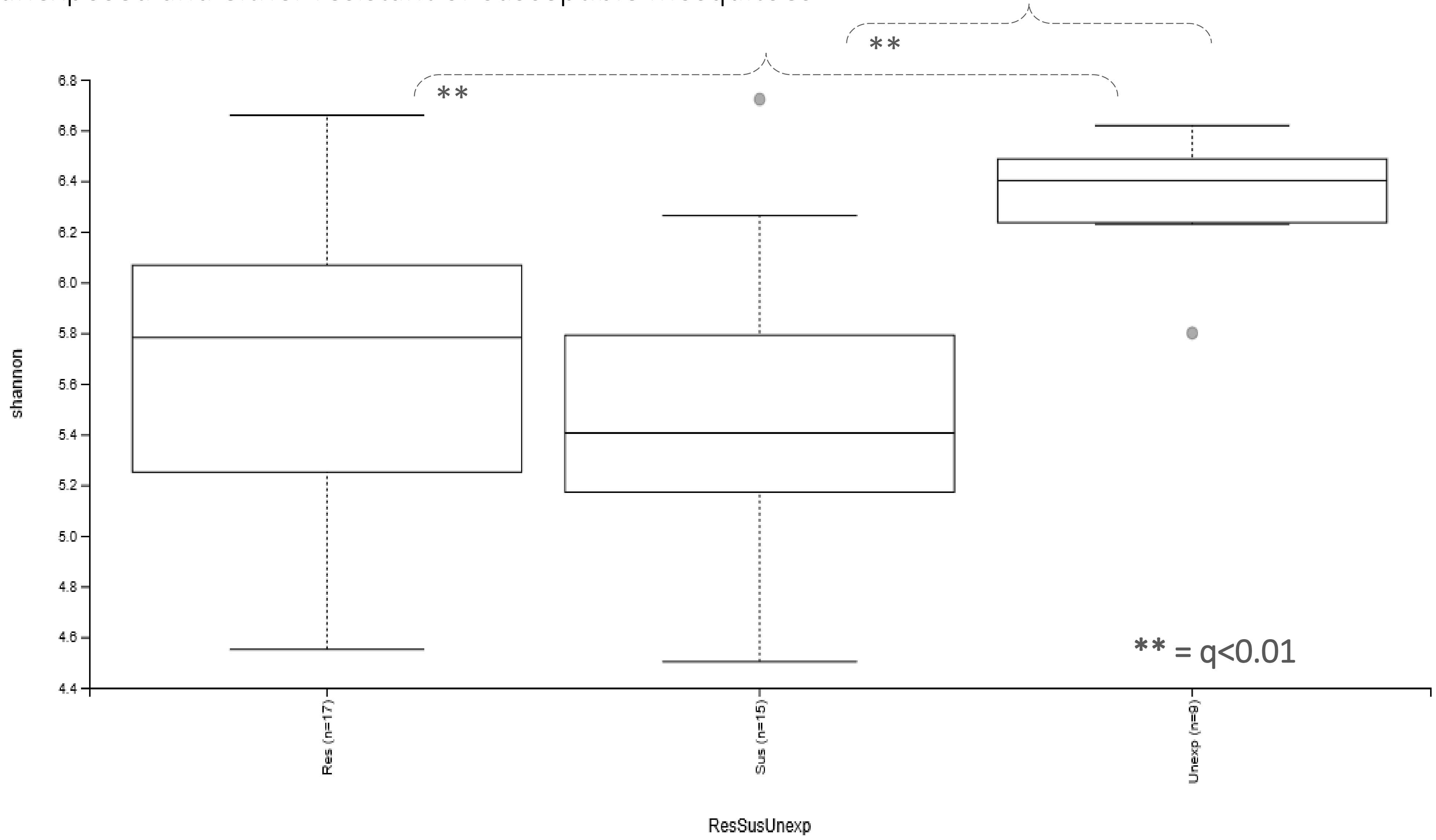

**Suppl. 5D. Insecticide exposure did not impact the internal bacterial abundance and evenness in *An. albimanus* larvae**. H = 2.7, p=0.25.

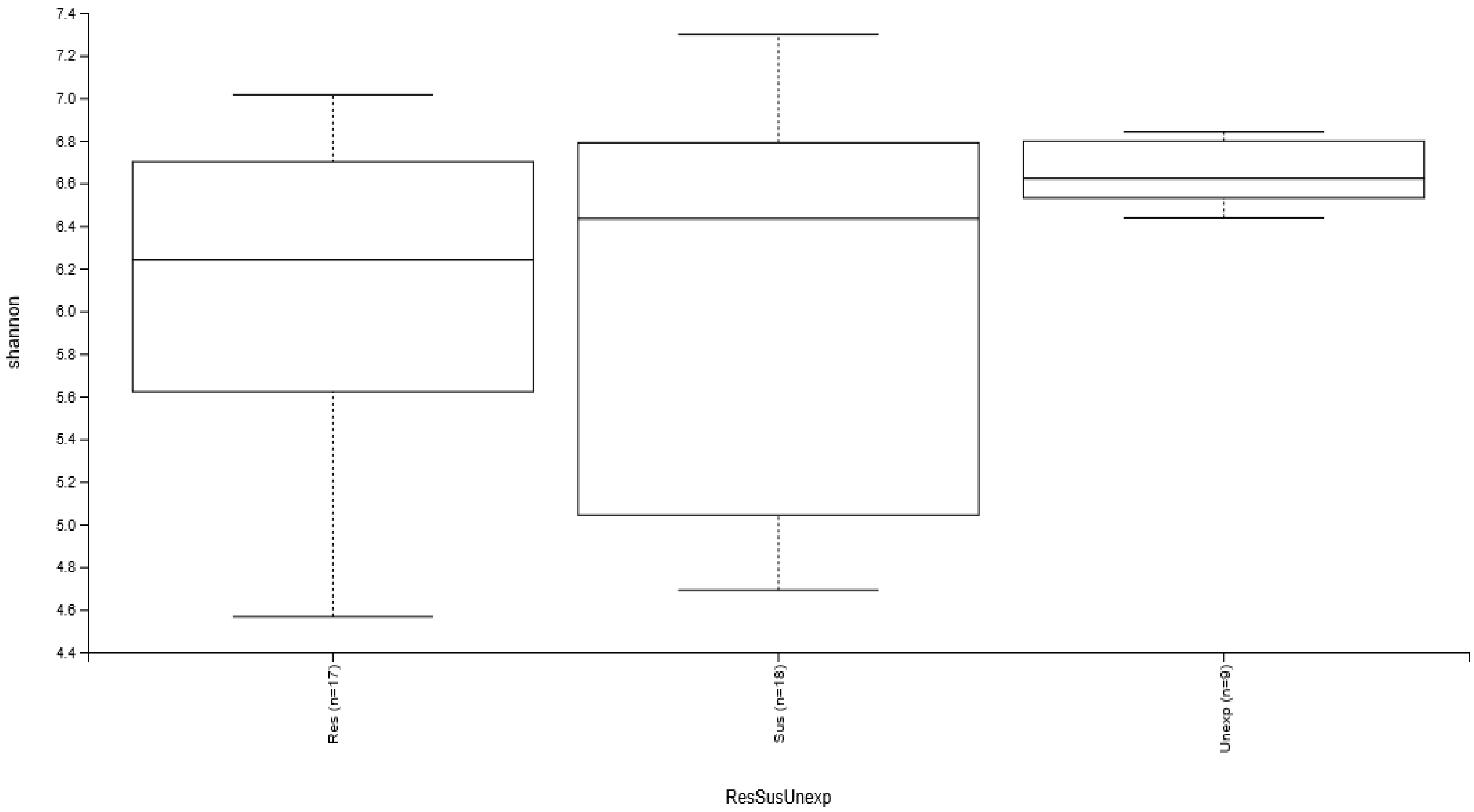

Suppl. 5E. Type of insecticide did not impact bacterial abundance and evenness on the cuticle surface of adult *An. albimanus*. H=3.8, p=0.28

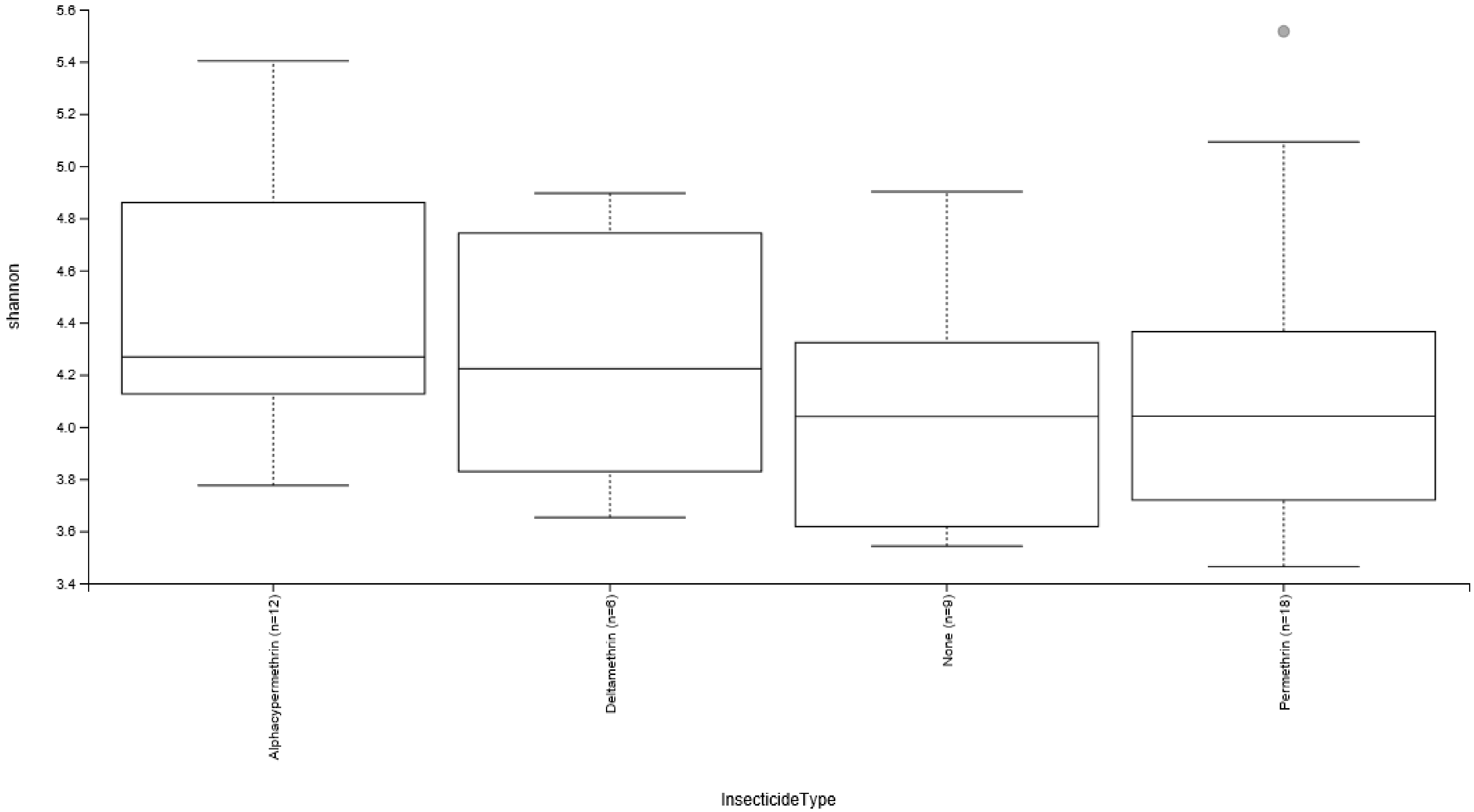

Suppl. 5F. Type of insecticide significantly impacted the abundance and evenness of internal bacteria in adult *An. albimanus*. H = 10.1, p=0.02. Deltamethrin-exposed adults represented the most abundant and even bacterial community

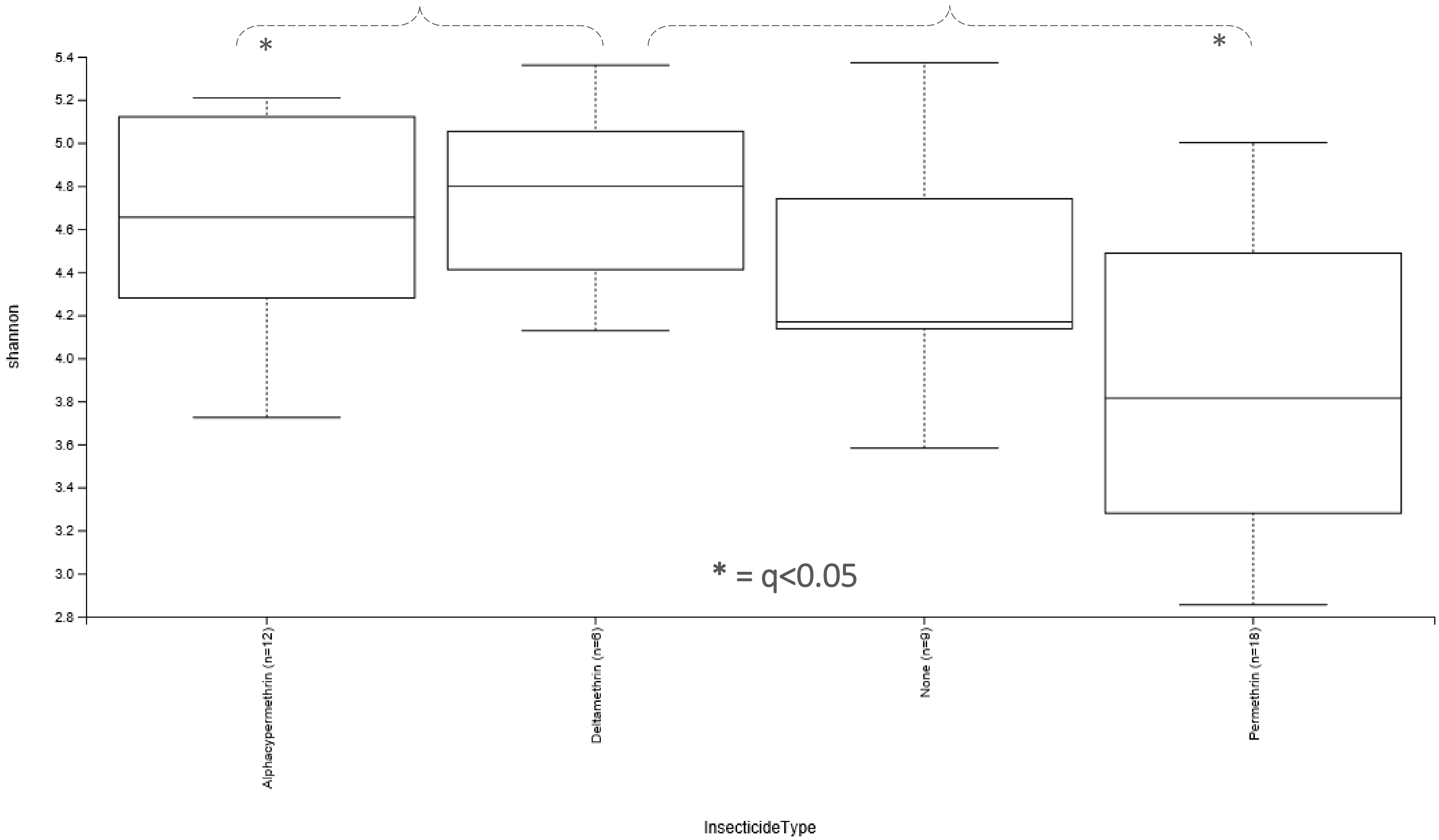

Suppl. 5G. Insecticide exposure did not significantly impact bacterial abundance and evenness on the cuticle surface of adult *An. albimanus*. H=0.73, p=0.69.

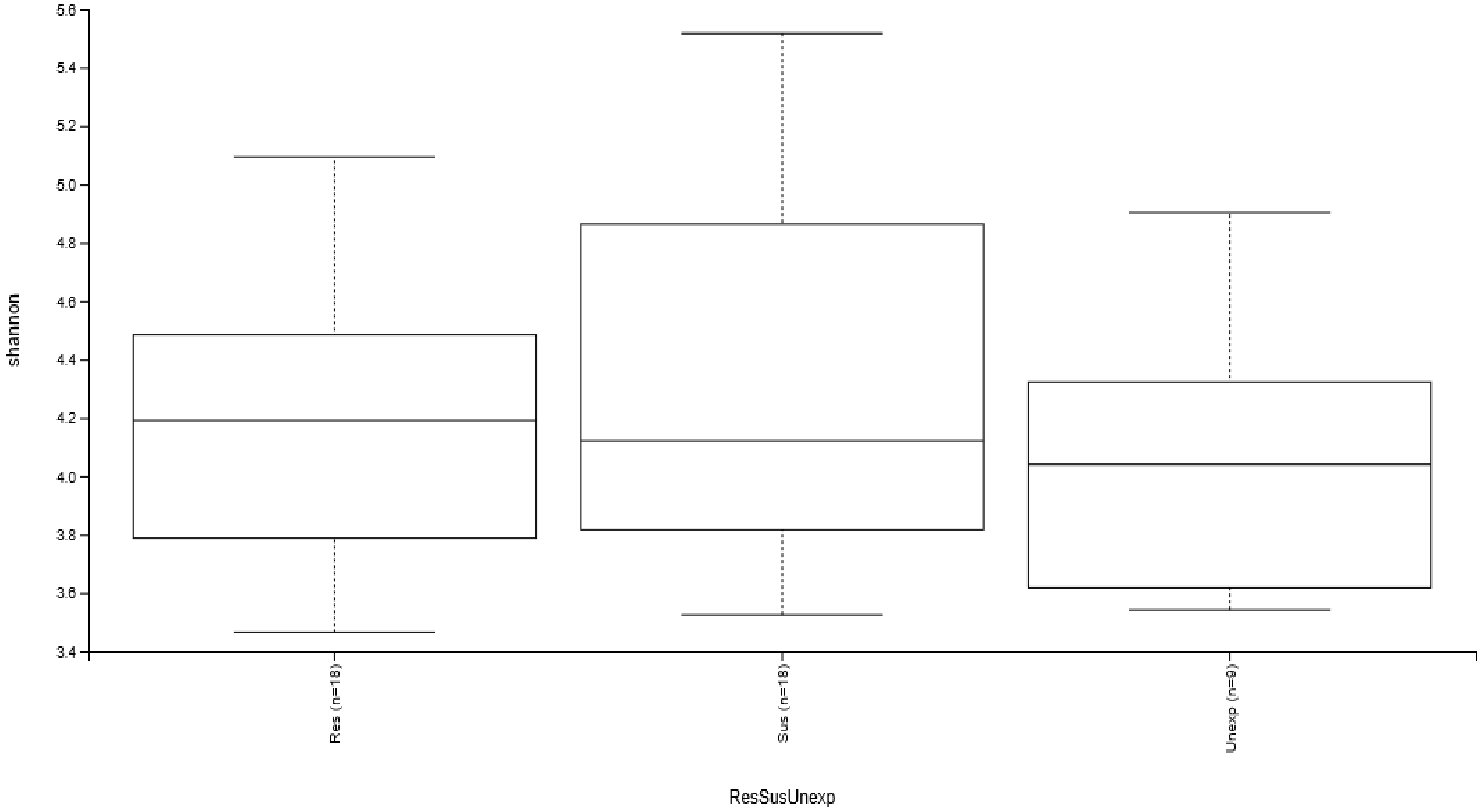

Suppl. 5H. Insecticide exposure did not significantly impact the abundance and evenness of internal bacteria in adult *An. albimanus*. H = 0.7, p=0.7

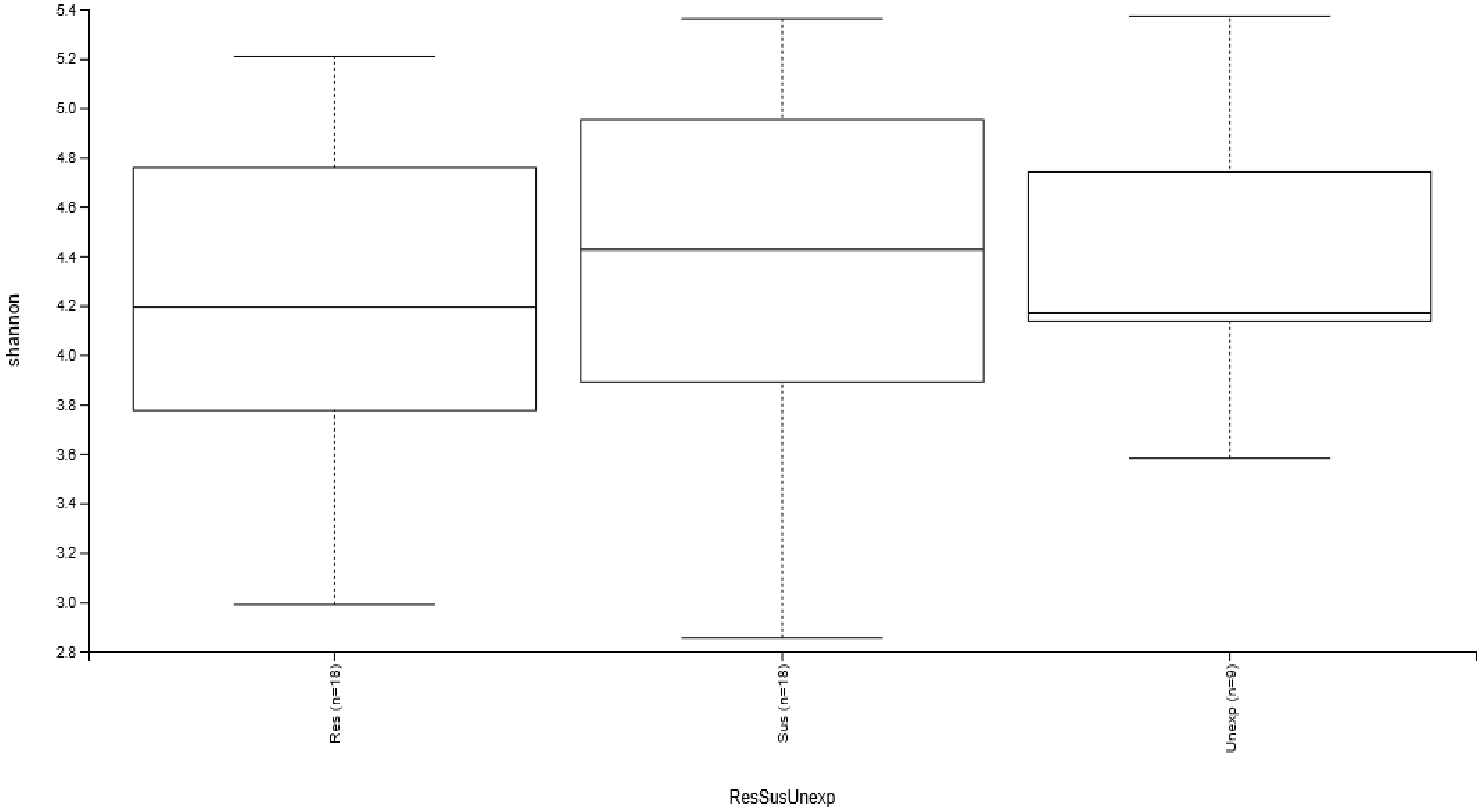
