## Supplementary material for "Pyrethroid exposure alters *Anopheles albimanus* microbiota and resistant mosquitoes harbor more insecticide-metabolizing bacteria": Suppl. 6

**Suppl. 6: Beta diversity comparisons show differential bacterial composition between pyrethroid-exposed and non-exposed *An. albimanus***

| | | | Internal microbiota | | | | Cuticle surface microbiota | | | |
|---|---|---|---|---|---|---|---|---|---|---|
| Stage | Group 1 | Group 2 | N | pseudo-F | p-value | q-value | N | pseudo-F | p-value | q-value |
| Adults | Resistant | Susceptible | 36 | 0.46 | 0.843 | 0.843 | 36 | 1.10 | 0.340 | 0.340 |
| | Resistant | Non-exposed | 27 | **11.80** | **0.001** | **0.002** | 27 | 1.71 | 0.102 | 0.306 |
| | Susceptible | Non-exposed | 27 | **9.44** | **0.001** | **0.002** | 27 | 1.27 | 0.240 | 0.340 |
| Larvae | Resistant | Susceptible | 35 | 1.06 | 0.335 | 0.335 | 32 | 1.04 | 0.340 | 0.340 |
| | Resistant | Non-exposed | 26 | 1.91 | 0.072 | 0.216 | 26 | **3.28** | **0.005** | **0.008** |
| | Susceptible | Non-exposed | 27 | 1.49 | 0.182 | 0.273 | 24 | **3.92** | **0.004** | **0.008** |

Pair-wise comparisons of beta diversity (Bray Curtis) between pyrethroid-exposed and non-exposed mosquitoes showed significant differences in adult internal, but not cuticle surface, bacterial composition. In contrast, this difference was evident in larval cuticle surface, but not internal, microbiota. There was no significant difference in bacterial composition between pyrethroid-resistant and –susceptible mosquitoes at both larval and adult stages. Comparisons were conducted using PERMANOVA (999 permutations) tests with Benjamini-Hochberg FDR correction (q-value). Significance is set to q-value (adjusted p-value) <0.05. N = number of mosquito pools included in the analysis; each pool comprised 3 mosquitoes.
