## Supplementary material for "Pyrethroid exposure alters *Anopheles albimanus* microbiota and resistant mosquitoes harbor more insecticide-metabolizing bacteria": Suppl. 7

**Suppl. 7: Beta diversity comparisons show differential bacterial composition between *Anopheles albimanus* that exposed or unexposed to alphacypermethrin and permethrin.**

| | | | | Internal microbiota | | | | Cuticle surface microbiota | | | |
|---|---|---|---|---|---|---|---|---|---|---|---|
| Stage | Insecticide | Group 1 | Group 2 | N | pseudo-F | p-value | q-value | N | pseudo-F | p-value | q-value |
| Adults | Alphacypermetrin | Resistant | Susceptible | 12 | 1.48 | 0.128 | 0.128 | 12 | 1.39 | 0.134 | 0.134 |
| | | Resistant | Unexposed | 15 | **6.05** | **0.001** | **0.002** | 15 | **3.10** | **0.014** | **0.021** |
| | | Susceptible | Unexposed | 15 | **5.16** | **0.001** | **0.002** | 15 | **2.59** | **0.008** | **0.021** |
| | Deltamethrin | Resistant | Susceptible | 6 | 1.01 | 0.421 | 0.421 | 6 | 0.53 | 1.000 | 1.000 |
| | | Resistant | Unexposed | 12 | 1.49 | 0.173 | 0.260 | 12 | 1.23 | 0.318 | 0.843 |
| | | Susceptible | Unexposed | 12 | 2.17 | 0.083 | 0.249 | 12 | 0.81 | 0.562 | 0.843 |
| | Permethrin | Resistant | Susceptible | 17 | 0.37 | 0.789 | 0.789 | 18 | 0.45 | 0.684 | 0.770 |
| | | Resistant | Unexposed | 18 | **12.29** | **0.001** | **0.002** | 18 | 0.85 | 0.595 | 0.770 |
| | | Susceptible | Unexposed | 17 | **9.64** | **0.001** | **0.002** | 18 | 0.66 | 0.770 | 0.770 |
| Larvae | Deltamethrin | Resistant | Susceptible | 17 | 0.41 | 0.810 | 0.810 | 16 | 0.76 | 0.538 | 0.538 |
| | | Resistant | Unexposed | 17 | 1.55 | 0.130 | 0.245 | 17 | 2.38 | 0.051 | 0.077 |
| | | Susceptible | Unexposed | 18 | 1.54 | 0.163 | 0.245 | 17 | 2.35 | 0.043 | 0.077 |
| | Permethrin | Resistant | Susceptible | 18 | 1.47 | 0.180 | 0.270 | 17 | 1.11 | 0.315 | 0.315 |
| | | Resistant | Unexposed | 18 | 1.99 | 0.053 | 0.159 | 18 | **2.90** | **0.018** | **0.030** |
| | | Susceptible | Unexposed | 18 | 1.22 | 0.280 | 0.280 | 17 | **2.95** | **0.020** | **0.030** |

Pair-wise comparisons of beta diversity (Bray Curtis) between insecticide-exposed (i.e. resistant and susceptible) and –unexposed mosquitoes show variable results when individual insecticides were considered. In adults, the bacterial composition of mosquitoes exposed (i.e. resistant and susceptible) to alphacypermethrin and permethrin differed significantly from those of unexposed mosquitoes. For both insecticides, this difference was evident in the internal microbiota, but only true for alphacypermethrin in the cuticle surface microbiota. Conversely, the bacterial composition on larval cuticle surface, but not the internal microbial niche, differed only between permethrin exposed and unexposed mosquitoes. No effect on bacterial composition was seen in deltamethrin-tested larvae or adults. Also, for each insecticide tested, in both larvae and adults, there was no significant difference in bacterial composition between insecticide resistant or susceptible mosquitoes. Comparisons were conducted using PERMANOVA (999 permutations) tests with Benjamini-Hochberg FDR correction (q-value). Significance is set to q-value (adjusted p-value) <0.05. N = number of mosquito pool included in the analysis; each pool comprised 3 mosquitoes.
