## Supplementary material for "Pyrethroid exposure alters *Anopheles albimanus* microbiota and resistant mosquitoes harbor more insecticide-metabolizing bacteria": Suppl. 8

**Suppl. 8: Bacterial composition significantly differ between mosquitoes exposed to different insecticides.**

a.

| Group 1 | Group 2 | N | pseudo-F | p-value | q-value |
|---|---|---|---|---|---|
| **Alphacypermethrin** | **Deltamethrin** | **67** | **17.42** | **0.001** | **0.002** |
| **Alphacypermethrin** | **None** | **60** | **10.26** | **0.001** | **0.002** |
| **Alphacypermethrin** | **Permethrin** | **96** | **10.10** | **0.001** | **0.002** |
| **Deltamethrin** | **None** | **79** | **3.13** | **0.017** | **0.026** |
| Deltamethrin | Permethrin | 115 | 2.11 | 0.073 | 0.073 |
| **None** | **Permethrin** | **108** | **2.43** | **0.028** | **0.034** |

b.

| | | | Internal microbiota | | | | Cuticle surface microbiota | | | |
|---|---|---|---|---|---|---|---|---|---|---|
| Stage | Group 1 | Group 2 | N | pseudo-F | p-value | q-value | N | pseudo-F | p-value | q-value |
| Adult | Alphacypermethrin | Deltamethrin | 18 | 1.43 | 0.222 | 0.222 | 18 | 1.28 | 0.25 | 0.375 |
| | **Alphacypermethrin** | **None** | **21** | **7.54** | **0.001** | **0.003** | **21** | **2.78** | **0.005** | **0.03** |
| | Alphacypermethrin | Permethrin | 30 | 2.13 | 0.057 | 0.068 | 30 | 1.75 | 0.123 | 0.369 |
| | **Deltamethrin** | **None** | **15** | **2.78** | **0.029** | **0.044** | 15 | 1.34 | 0.2 | 0.375 |
| | **Deltamethrin** | **Permethrin** | **24** | **5.65** | **0.002** | **0.004** | 24 | 0.83 | 0.494 | 0.593 |
| | **None** | **Permethrin** | **27** | **16.41** | **0.001** | **0.003** | 27 | 0.74 | 0.619 | 0.619 |
| Larvae | **Deltamethrin** | **None** | **26** | 1.91 | 0.075 | 0.225 | 23 | **3.26** | **0.016** | **0.024** |
| | Deltamethrin | Permethrin | 35 | 0.86 | 0.514 | 0.514 | 32 | 1.03 | 0.365 | 0.365 |
| | **None** | **Permethrin** | **27** | 1.44 | 0.173 | 0.26 | 27 | **3.83** | **0.002** | **0.006** |

Pair-wise comparisons of beta diversity (Bray Curtis) between groups of *An. albimanus* that were exposed to alphacypermethrin, deltamethrin or permethrin overall (a) and by developmental stage (b) show that bacterial composition differ significantly between mosquitoes exposed to different insecticides. Comparisons were conducted using PERMANOVA (999 permutations) tests with Benjamini-Hochberg FDR correction (q-value). Significance is set to q-value (adjusted p-value) <0.05. N = number of mosquito pool included in the analysis; each pool comprised 3 mosquitoes.
